# Mitochondria supply ATP to the ER through a mechanism antagonized by cytosolic Ca^2+^

**DOI:** 10.1101/591685

**Authors:** Jing Yong, Helmut Bischof, Marina Siirin, Anne Murphy, Roland Malli, Randal J. Kaufman

## Abstract

The endoplasmic reticulum (**ER**) imports ATP and uses energy from ATP hydrolysis for protein folding and trafficking. However, little is known about this vital ATP transport process across the ER membrane. Here, using three commonly used cell lines (CHO, INS1 and HeLa), we report that ATP enters the ER lumen through a cytosolic Ca^2+^-antagonized mechanism, or *CaATiER* (**Ca^2+^-A**ntagonized **T**ransport into **ER**) mechanism for brevity. Significantly, we observed that a Ca^2+^ gradient across the ER membrane is necessary for ATP transport into the ER. Therefore Ca^2+^ signaling in the cytosol is inevitably coupled with ATP supply to the ER. We propose that under physiological conditions, cytosolic Ca^2+^ inhibits ATP import into the ER lumen to limit ER ATP consumption. Furthermore, the ATP level in the ER is readily depleted by oxidative phosphorylation (OxPhos) inhibitors, and that ER protein misfolding increases ATP trafficking from mitochondria into the ER. These findings suggest that ATP usage in the ER may increase mitochondrial OxPhos while decreasing glycolysis, i.e., an “*anti-Warburg*” effect.

**Significance Statement:** We report that ATP enters the ER lumen through an AXER-dependent, cytosolic Ca^2+^-antagonized mechanism, or *CaATiER* (Ca^2+^-Antagonized Transport into ER) mechanism. In addition, our findings suggest that ATP usage in the ER may render an “anti-Warburg” effect by increasing ATP regeneration from mitochondrial OxPhos while decreasing the portion of ATP regeneration from glycolysis.

---

Energy supply is a fundamental requirement for all cells to perform their biochemical functions. Universally, ATP is the single most important energy-supplying molecule in every form of life. ATP regeneration from ADP takes place in mitochondria mainly through OxPhos, and in the cytosol through glycolysis. Despite its heavy demand for ATP to facilitate protein folding and trafficking, the ER is not known to possess an independent ATP regeneration machinery. A protein transporter, “*ER-ANT*” is involved in the ATP translocation across the ER membrane, of which the biochemical properties are analogous to the mitochondrial **A**denosine **N**ucleotide **T**ransporter (**ANT**) (1, 2). Other than that, little is known about how ATP gets into the ER in a living cell or whether/how ATP consumption is regulated in the ER lumen. For example, only one report described the genetic identification of an ATP transporter *ER-ANT1* in *Arabidopsis* and its deletion caused a disastrous plant phenotype, characterized by drastic growth retardation and impaired root and seed development (3). The mammalian equivalent of *ER-ANT1* had remained elusive until a recent publication identified SLC35B1 as the putative mammalian ER ATP transporter (4).

On the other hand, ATP in the ER is essential to support protein chaperone functions for protein folding, such as BiP/GRP78, and trafficking (5–9). In fact, the level of ATP determines which proteins are able to transit to the cell surface (5, 6). Although the level of ER ATP is suggested to impact protein secretion, this has not been demonstrated, nor have the factors that regulate ATP levels in the ER been elucidated. With two recently characterized genetically encoded ATP reporter proteins targeted to select intracellular organelles, namely the mitochondrial localized *AT1.03* and the ER localized *ERAT4* probes (10, 11), we studied the ATP status within the ER organelles in intact cells, in parallel with another study of the ATP status in the mitochondria matrix (12). Here, we monitored real time changes in ATP levels inside the ER lumen in response to well-characterized OxPhos and/or glycolysis inhibitors in living Chinese hamster ovary (**CHO**), rat insulinoma INS1 and human Hela cells, at the single cell level using an ERAT-based FRET assay. In addition, we monitored the change in ER ATP upon Ca^2+^ release from the ER, and further evaluated the ER ATP status in response to varying cytosolic Ca^2+^ concentrations. Our findings lead us to propose that cytosolic Ca^2+^ attenuates mitochondrial-driven ATP transport into the ER lumen through a *CaATiER* (**Ca^2+^-A**ntagonized **T**ransport into **ER**) mechanism. This model was further validated by knocking-down *Slc35b1* in human HeLa cells (4), and under conditions of ER protein misfolding in CHO cells.

## Results

### ER ATP comes from Mitochondrial OxPhos in CHO cells

Traditional ATP analytical methods based on biochemical or enzymatic assays inevitably require ATP liberation from endogenous compartments, and do not reflect compartment-specific ATP dynamics. Nevertheless, there is ample evidence supporting that differential ATP levels exist in membrane-bound organelles that use independent regulatory mechanisms in a compartment-specific manner (10–13). To detect ATP levels in the ER lumen *in vivo*, we expressed an ER-localized ATP sensor ERAT (ERAT4.01^N7Q^) in H9 CHO cells engineered to induce mRNA expression of human clotting factor VIII (**F8**), encoding a protein which misfolds in the ER lumen, upon histone deacetylase inhibition (14, 15). Confocal analysis of ERAT fluorescence (**Fig. 1A**, green) revealed nearly complete co-localization with the ER marker, ER-Tracker™ Red (**Fig. 1A**, red), as well as with the endogenous ER-resident protein PDIA6 detected by immunofluorescence (**Sppl. Fig. S1A**). Induction of F8 by SAHA treatment for 20 hrs did not change the ER localization of the ERAT reporter (**Fig. 1B and Sppl Fig. S1B**). Another protein ER marker, ER-RFP, also shows nearly complete co-localization with ERAT fluorescence although ER-RFP formed aggregates (16) upon SAHA induction (**Sppl Fig. S2 A and B**, with RFP aggregates indicated with yellow arrow heads). As there is no known ATP regeneration machinery in the ER, and intracellular ATP regeneration from ADP takes place in mitochondria through OxPhos, and in the cytosol through glycolysis, it is of key importance to determine whether the ER ATP status depends on OxPhos and/or glycolysis (illustrated in **Fig. 1C**). For this purpose, we analyzed ER ATP levels after OxPhos inhibition or glycolysis inhibition by monitoring the FRET signal through flow-cytometry, since this technique provides robust population statistics and circumvents photo-bleaching of the fluorescent probes. A detailed description of the experimental procedures using the H9-D2 cell line specifically engineered for the flow-based FRET assay can be found in the “*Supplemental Materials*” section.

**Figure 1.**
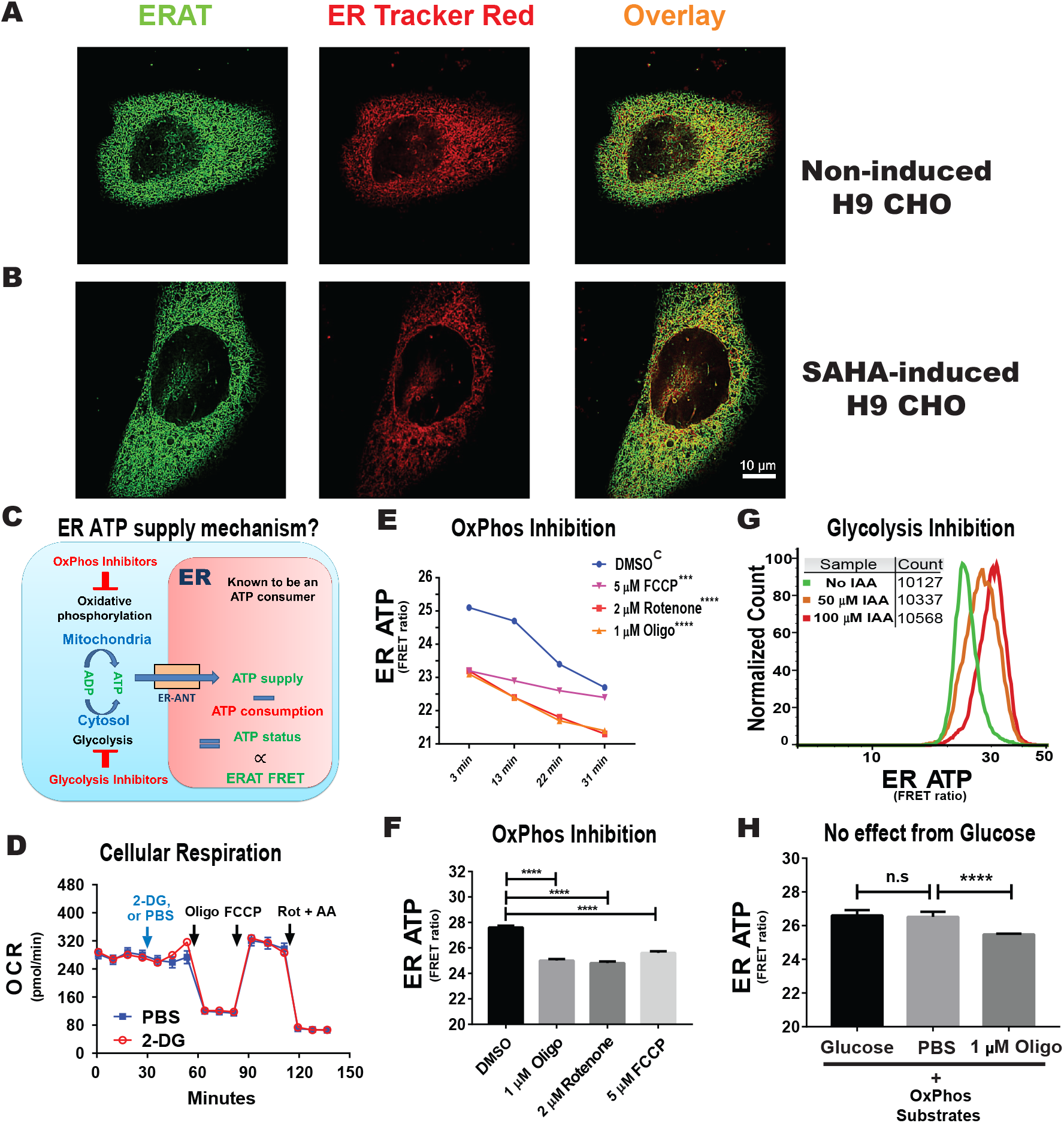
ER ATP homeostasis is maintained by oxidative phosphorylation. **A.** Confocal microscopy confirms ER localization of the ER ATP FRET reporter, ERAT in H9-D2 CHO cells. A representative confocal micrograph shows a high degree of co-localization of ERAT fluorescence in green with ER-Tracker™ Red, in red. **B.** Increased F8 expression in H9-D2 CHO cells by SAHA treatment does not alter ERAT’s co-localization with ER-Tracker™ Red. (Scale bar: 10 μm) **C.** A cartoon model depicting the ET ATP monitoring system. Importantly, the ER is an obligate ATP consumer as there is no known ATP regeneration system inside the ER lumen. ERAT reports steady state ATP levels inside the ER, which is a dynamic balance between ATP supply and ATP consumption. **D.** 2-DG (20 mM) has no effect on cellular respiration. Oxygen consumption rates (OCR, in pMole/min) in H9 CHO cells were measured by an *XF-24* platform (Seahorse BioScience), using serial injections of the compounds; first PBS or 2-DG followed sequentially by 1 μM oligomycin (Oligo), 1 μM FCCP and 1 μM rotenone together with 10 μM antimycin A (Rot + AA). **E.** The effect of OxPhos inhibitors (oligomycin, rotenone and FCCP at the indicated concentrations) on ER ATP was examined by flow cytometry at four time points as indicated. The same tubes of suspended H9 CHO cells were repeatedly sampled after OxPhos inhibition. Statistical significance values for curve comparisons by two-way ANOVA are labeled for individual inhibitors versus the “DMSO” control group marked by an upper case “C”. **F.** Administration of OxPhos inhibitors for 15 min, represented by oligomycin, rotenone and FCCP, rapidly reduces ER ATP in H9 CHO cells. DMSO was included as solvent control for comparison. Statistical significance values are labeled for individual comparisons. **G.** Administration of IAA, a glycolysis inhibitor, increases ER ATP levels (detected by flow cytometry) in H9 CHO cells. Experiments were repeated four times, and a representative result is shown. Specifically, histograms of ER ATP levels for individual cells at two IAA concentrations (50 μM and 100 μM, treated for 2 hrs) are shown with the y-axis representing normalized cell counts. More than ten thousand cells were sampled for every treatment condition, as indicated on the graph. **H.** Glucose supplementation (5 mM) to DMEM medium with only OxPhos substrates (with 1 mM sodium pyruvate supplemented) does not affect ER ATP levels. H9 CHO cells stably transfected with the ERAT reporter were incubated in the serum-free modified DMEM medium, which contains substrates (i.e. pyruvate plus L-Ala-Gln) to support only OxPhos, for 1 hr at 37 °C in a cell culture incubator, before flow analysis for ER ATP status. For the “1 μM Oligo” group, oligomycin was added to cells for the last 20 min of incubation in glucose-free medium. *Two-way ANOVA* was applied for statistical analysis of geometric means of fluorescence intensity (gMFI), with significance levels expressed as: *n.s*: - not significant; * - p≤ 0.05; **- p≤ 0.01; ***- p< 0.001; **** - p< 0.0001. The same statistical analysis was applied in the following figures.

In addition, as inhibition of glycolysis may have adverse effects on OxPhos, we tested whether 2-deoxyglucose (**2-DG**) administration reduces mitochondrial respiration by measuring the cellular oxygen consumption rate (**OCR**) in CHO-H9 cells using a *Seahorse XF24*^®^ respirometry platform. Importantly, with glucose and pyruvate supplemented in the medium to support mitochondrial respiration, 2-DG administration did not negatively affect the OCR, under both coupled and uncoupled respiration conditions (**Fig. 1D** and **Sppl. Fig. S3A**), suggesting that glycolysis inhibition by 2-DG does not acutely impair OxPhos in CHO cells. Interestingly, oligomycin (**Oligo**) treatment immediately increased the extracellular acidification rate (**ECAR**) in H9 CHO cells, reflecting glycolysis upregulation to compensate for ATP shortage due to OxPhos inhibition, while 2-DG treatment inhibited this compensation (**Sppl. Fig. S3B**).

We subsequently tested three potent OxPhos inhibitors, oligomycin (targeting ATP synthase, also known as complex V), rotenone (targeting complex I), and FCCP (a protonophore that depolarizes the mitochondrial inner membrane to uncouple ATP synthase). All OxPhos inhibitors significantly decreased ER ATP levels immediately after administration, by analysis of ER ATP kinetics with time (**Fig. 1E**) or by analysis at a single time point (**Fig. 1F**). Importantly, the bio-energetic inhibitors used had no effect on ER localization of the ERAT probe, as shown by confocal microscopy in H9 CHO cells co-transfected with a cytosolic RFP-based reporter, Cyto TagRFP (16) (**Sppl. Fig. S4 A – F**), and was further reflected by no change of the *Pearson* correlation coefficient under these conditions (**Sppl Fig. S4G**). As predicted by the chemical law of mass action, the decrease in ER ATP supply upon OxPhos inhibition should reduce ER ATP consumption. Furthermore, if we assume that the three OxPhos inhibitors do not stimulate ATP usage in the ER, the results (**Fig. 1E and F**) indicate that the steady-state ER ATP supply is heavily dependent on OxPhos. Since the FRET signal generated from the ERAT reporter is ratiometric and does not provide a direct calibration corresponding to ATP concentration in the ER in live cells, we attempted to determine the effect of complete inhibition of cellular ATP regeneration thereby providing the lowest detection range of the ERAT reporter by treating cells for prolonged time (2 hrs) with 2-DG, followed by oligomycin treatment (**Sppl. Fig. S5 A** and **B**). This condition successfully identified the lower limit of the dynamic range for the ERAT reporter when the ratiometric FRET/CFP signal no longer decreases.

In contrast to OxPhos inhibition, the interpretation of effects from glycolysis inhibition is rather complicated, as there is no pure “glycolysis inhibitor” *per se*, e.g. 2-DG application consumes cytosolic ATP and may even increase intracellular ADP levels as a substrate to boost mitochondrial ATP production through OxPhos (**Fig. 1D** and **Sppl. Fig. S3A**). With this caveat in mind, it was particularly intriguing to observe that incubation with iodoacetamide (**IAA**), an inhibitor of glyceraldehyde phosphate dehydrogenase (**GAPDH**), for 2 hrs increased steady state ER ATP levels in H9 CHO cells, in a dose-dependent manner (illustrated in **Fig. 1G**). In an effort to further measure the extent by which glycolysis-derived ATP contributes to the ER ATP pool, we found that glucose supplementation had no effect on ER ATP levels, in a defined medium with substrates supporting only OxPhos (**Fig. 1H**), while oligomycin strongly decreased ER ATP under these conditions (p< 0.0001, with representative traces of raw FRET ratios shown in **Sppl. Fig. S7A**). In contrast, glucose supplementation significantly increased cellular ATP measured by luciferase assays under the same medium conditions (**Sppl. Fig. S6A**), and 2-DG decreased cellular ATP, while oligomycin only had a minor effect on cellular ATP compared to 2-DG (**Sppl. Fig. S6B**). These observations are consistent with the notion that CHO cells are highly dependent on glycolysis to generate ATP. Furthermore, 2-DG immediately decreased the cytosolic ATP/ADP ratio measured by flow-cytometry using the ratiometric *PercevalHR* probe (**Sppl. Fig. S6C**), while OxPhos inhibition by oligomycin did not significantly alter this ratio (**Sppl. Fig. S6D**), as previously shown by others (10, 17). These results provide strong evidence supporting the notion that distinct ATP supplying and regulatory mechanisms exist for subcellular compartments within a cell, as we reported (12). Furthermore, a third widely used glycolysis inhibitor, 3-bromopyruvate (**3-BrP**), produced a similar ATP increase in the ER, in a dose dependent manner (**Sppl. Fig. S7B**). At the same time, the kinetics of the ER ATP increase by 3-BrP was distinct from that by IAA, suggesting compound-specific off-target effects by the “glycolysis inhibitors” (**Sppl. Fig. S7C**). During the flow cytometry data analysis, we noticed a reproducible decrease in steady-state ER ATP levels in the control group of H9-D2 CHO cells, especially after 30 mins of cell suspension (examples as shown in **Fig. 1E** and **Fig. 2 E&F**). We suspect this baseline ATP drift could be caused by the flow cytometry procedure, as extracellular matrix-attachment is crucial for optimal mitochondrial respiration (18, 19). We acknowledge this feature as a disadvantage for flow-based ER ATP assay, and therefore have also included fluorescent microscopy to validate our findings. Finally, the observations made in H9 CHO cells were completely reproduced in CHO cells that do not express human F8 (20), i.e. oligomycin or rotenone administration reduced steady state ER ATP levels (**Sppl. Fig. S7D**, indicated by red and orange curves). In contrast, inhibitors of glycolysis, 2-DG or IAA, immediately increased ER ATP levels (**Sppl. Fig. S7D**, indicated by dark green and light green curves).

**Figure 2.**
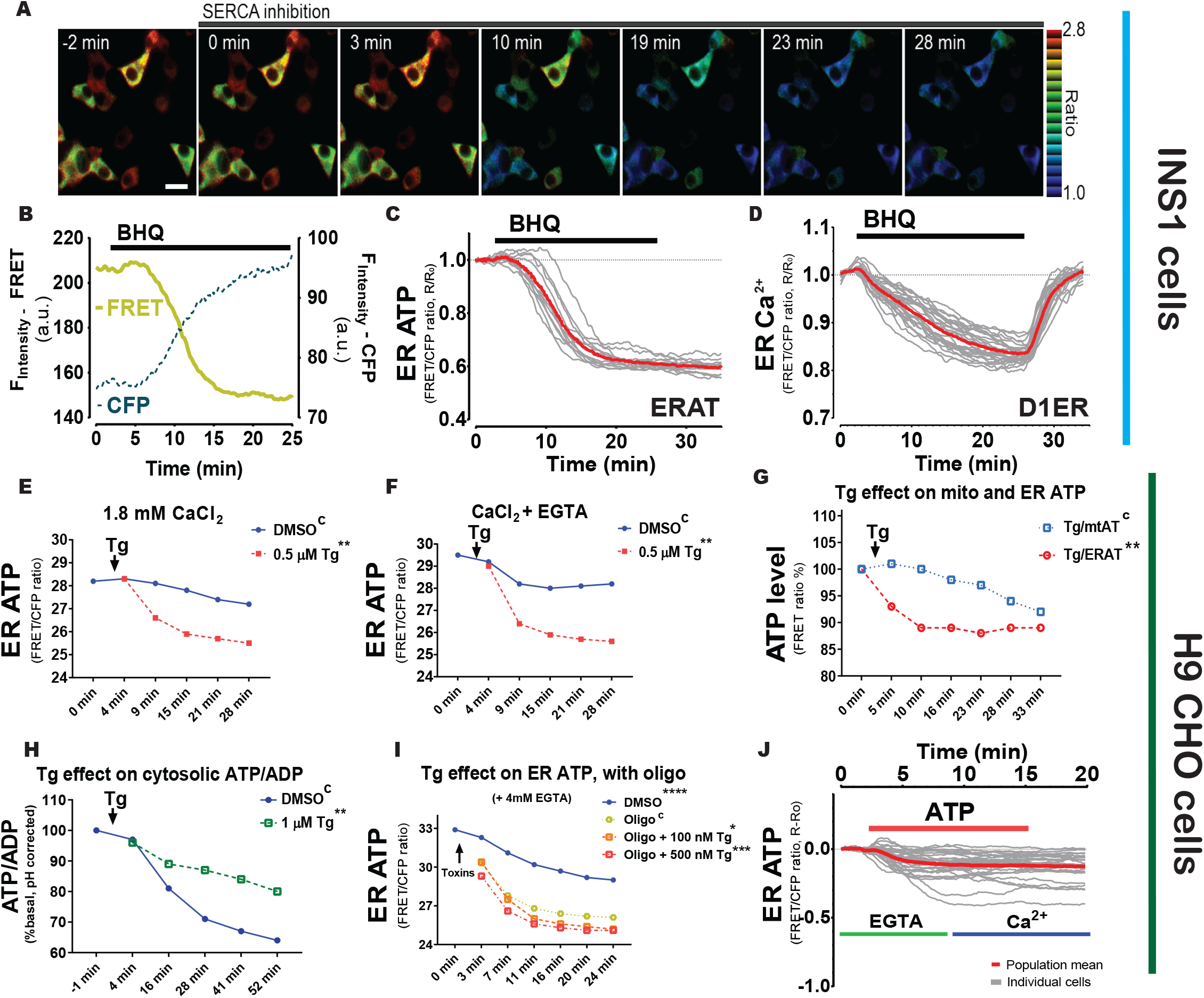
A Ca^2+^ gradient across the ER membrane is required to maintain ER ATP homeostasis. **A.** SERCA inhibition reduces ER ATP levels in adherent INS1 cells. BHQ (15 μM) was added at 0 min as indicated. Scale bar represents 10 μm. **B**. A representative trace of the FRET signal overlaid with the CFP fluorescence intensity is shown for illustration purposes in an adherent INS1 cell. **C**. BHQ reduces ER ATP. Y-axis represents the ratio between FRET intensity versus CFP fluorescence intensity. The average ratio is shown in red bold solid line in the graph, with individual INS1 cells shown in gray (n=13 cells from 3 experiments). **D**. Quantification of the effect of BHQ on ER Ca^2+^ levels in individual INS1 cells. The average ratio from the Ca^2+^ reporter (D1ER) is shown by a red solid line, with individual cells shown in gray (n=25 cells from 3 experiments). Notably, BHQ withdrawal at 25 min by perfusion causes Ca^2+^ refilling in the ER by SERCA re-activation. **E**. Thapsigargin (**Tg**, 0.5 μM added at 0 min) reduces ER ATP significantly. Flow analysis was performed on H9 CHO cells containing the ERAT reporter and with 1.82 mM CaCl_2_ in the culture medium. Statistical significance is labeled in the figure compared to the “*DMSO*” group as control, indicated with an upper case “C”. **F**. In the same experiment as shown in panel E, when extra-cellular Ca^2+^ in the medium was chelated by adding 4 mM EGTA, Tg shows the same effects as shown in panel E, i.e., Tg decreases ER ATP significantly. **G**. In response to Tg (0.25 μM final concentration, added at 0 min), the ER ATP concentration in H9 CHO decreases significantly faster than the decrease in mitochondrial ATP. ER ATP levels were measured by the ERAT probe and mitochondrial ATP was measured by the *mtAT 1.03* probe. ATP levels in the two compartments were measured simultaneously in two sets of H9 CHO cells and shared the same readout as the FRET versus CFP ratio of the probes (F_530_/F_445_). The ATP levels measured by FRET ratios were further used to calculate the Tg-induced ATP decrease standardized to the respective DMSO-treated control groups, and ATP levels are expressed as percentage of the “*DMSO*” group (% of DMSO). **H**. Tg treatment (1 μM final concentration) increases the cytosolic ATP to ADP ratio in H9 CHO cells monitored by the *PercevalHR* reporter. Raw F_488_/F_405_ values from *PercevalHR* were corrected for the cytosolic cpYFP signal (F_488_/F_405_) in a different set of H9 CHO cells to compensate for the fluorescence change from pH change in the cytosol in response to Tg or DMSO (17). **I**. In the presence of oligomycin, Tg-induced ER Ca^2+^ depletion decreases ER ATP more quickly compared to oligomycin (1 μM) alone, and the effect is dependent on the Tg concentration, i.e., 500 nM Tg induced a faster decrease than 100 nM Tg. **J**. Addition of extracellular ATP (100 μM) decreases the ER ATP concentration in intact H9 CHO cells, in the presence (green line) or absence (blue line) of Ca^2+^ chelation by EGTA. In all panels, the ER ATP concentration was measured by the *ERAT4* (N7Q) probe and the y-axis indicates the FRET ratio derived by dividing fluorescence intensity at 530 nm (F_530_) by that at 445 nm (F_445_), with a 405 nm laser excitation, i.e., FRET ratio = F_530_/F_445_. Statistical significance is shown for the indicated concentrations, compared to the control group.

To conclude, OxPhos-derived ATP constitutes the major ATP supplying source for the ER compartment, while it is not possible to conclude for the ER ATP dependence on glycolysis-generated ATP, as glycolysis inhibition inevitably leads to unknown effects on ER ATP usage (**Fig. 1C**) that may produce a net increase in ER ATP.

### ER ATP import requires a Ca^2+^ gradient across the ER membrane

We propose that protein misfolding in the ER disrupts ER ATP homeostasis. Consistent with this notion, application of the reversible SERCA inhibitor, 2,5-di-(tert-butyl)-1,4-hydroquinone (**BHQ**), caused a rapid decrease in ER ATP in rat insulinoma INS1 cells (**Fig. 2A**). INS1 cells were selected because they secrete large amounts of another protein, proinsulin and insulin. As we reported previously (11), due to reduced ATP levels in the ER lumen, the FRET fluorescence intensity from the ERAT probe decreased while the corresponding CFP fluorescence intensity increased simultaneously (**Fig. 2B**). The average of FRET over CFP ratios in a population of INS1 cells decreased and reached a plateau after 10 min post BHQ perfusion (**Fig. 2C**), and simultaneous measurement of ER Ca^2+^ by the D1ER probe (21) in another set of INS1 cells confirmed the specific BHQ effect on decreasing the ER Ca^2+^ concentration (**Fig. 2D**). We observed that ER ATP levels did not completely recover after BHQ removal, while in the same time frame, the ER Ca^2+^ concentration returned to its basal level (compare **Fig. 2C** to **2D**). It is particularly interesting that the ER ATP levels took longer to recover when we monitored the INS1 cells for up to 4 hours after BHQ withdrawal (**Sppl. Fig. S8A** and **B**). Furthermore, the recovered ER ATP levels in the same INS1 cells are responsive to a second BHQ stimulation in an extent proportional to the levels of ER ATP recovery (compare **Sppl. Fig. S8B** to **S8C**). These phenomena suggest that the Ca^2+^-responsiveness of the putative ER ATP transporter has a “memory” that lasts longer than the ER Ca^2+^ perturbation. In a similar manner, SERCA inhibition by thapsigargin (**Tg**), an irreversible SERCA inhibitor, rapidly decreased ER ATP levels in H9 CHO cells, reflected by a decrease in the ratio of FRET-to-CFP fluorescence intensity measured by flow cytometery (**Fig. 2E**). SERCA inhibition by Tg decreased ER ATP, whether the extracellular medium contained 1.8 mM Ca^2+^ (alpha MEM medium) (**Fig. 2E**) or no extracellular Ca^2+^ (by 2 mM EGTA chelation, **Fig. 2F**). The decrease in ER ATP by Tg correlated well with ER Ca^2+^ depletion as monitored by the GEM-CEPIA1er probe (22) (**Sppl. Fig. S9A** and **S9B**). Mechanistically, Tg disrupts the ER to cytosol Ca^2+^ gradient by inhibiting SERCA, a Ca^2+^ pump that shuttles 2 moles of Ca^2+^ from the cytosol into the ER lumen at the expense of 1 mole of ATP (23). Since maintaining low cytosolic Ca^2+^ concentrations (~100 nM under physiological conditions) also requires the action of plasma membrane Ca^2+^ pumps (**PMCA**) that consume ATP, it is possible that the decrease in ER ATP upon Tg treatment reflects mitochondrial ATP depletion due to excessive ATP usage in the cytosol. This possibility was tested by simultaneously measuring ER ATP in two sets of H9 CHO cells expressing either the ERAT probe (**Sppl. Fig. S9C**) or the mitochondrial ATP probe *mtAT*(10) (**Sppl. Fig. S9D**). Comparison of the decrease in relative ATP levels triggered by Tg revealed that the decrease in ER ATP preceded the decrease in mitochondrial ATP (**Fig. 2G**), which refutes the notion that Tg decreases ER ATP as a consequence of decreasing mitochondrial ATP. A third SERCA inhibitor, cyclopiazonic acid (**CPA**), induced a similar decrease in ER ATP at a dosage fifty times higher than Tg, presumably due to its low potency for inhibiting SERCA (**Sppl. Fig. S9A**).

In contrast, when the cytosolic ATP/ADP status was monitored by a structurally unrelated reporter, *PercevalHR* (17), Tg treatment attenuated the decrease in cytosolic ATP/ADP ratio in H9 CHO cells (**Fig. 2H**), which may be explained the effect of Tg on reducing cytosolic ATP usage by SERCA or by reducing ER ATP supply (see below in **Fig. 3G**). Tg-mediated depletion of the ER Ca^2+^ store did not appear to increase ATP usage in the ER since oligomycin treatment together with Tg reduced ER ATP only modestly, albeit significantly, compared to oligomycin alone (**Fig. 2I**). In addition, we investigated if Ca^2+^ release mediated through IP_3_R channel opening can affect ER ATP status, using a perfusion system and FRET-based fluorescence microscopy. Extracellular ATP is a well-known physiological IP3-generating agonist through its engagement with purinergic receptors ubiquitously expressed on CHO cells. Indeed, exogenously supplemented ATP in the perfusion buffer noticeably decreased the ER ATP pool (**Fig. 2J**). Furthermore, the effect of exogenous ATP was neither reversed by addition of extracellular Ca^2+^ (indicated by the blue bar in **Fig. 2J**) or washout of the agonist, again demonstrating that Ca^2+^ mobilization lowers ER ATP levels in a sustained manner and supporting a “memory” effect of the putative Ca^2+^ responsive ER ATP transporter. To summarize, ATP import into the ER apparently requires a Ca^2+^ gradient across the ER membrane.

**Figure 3.**
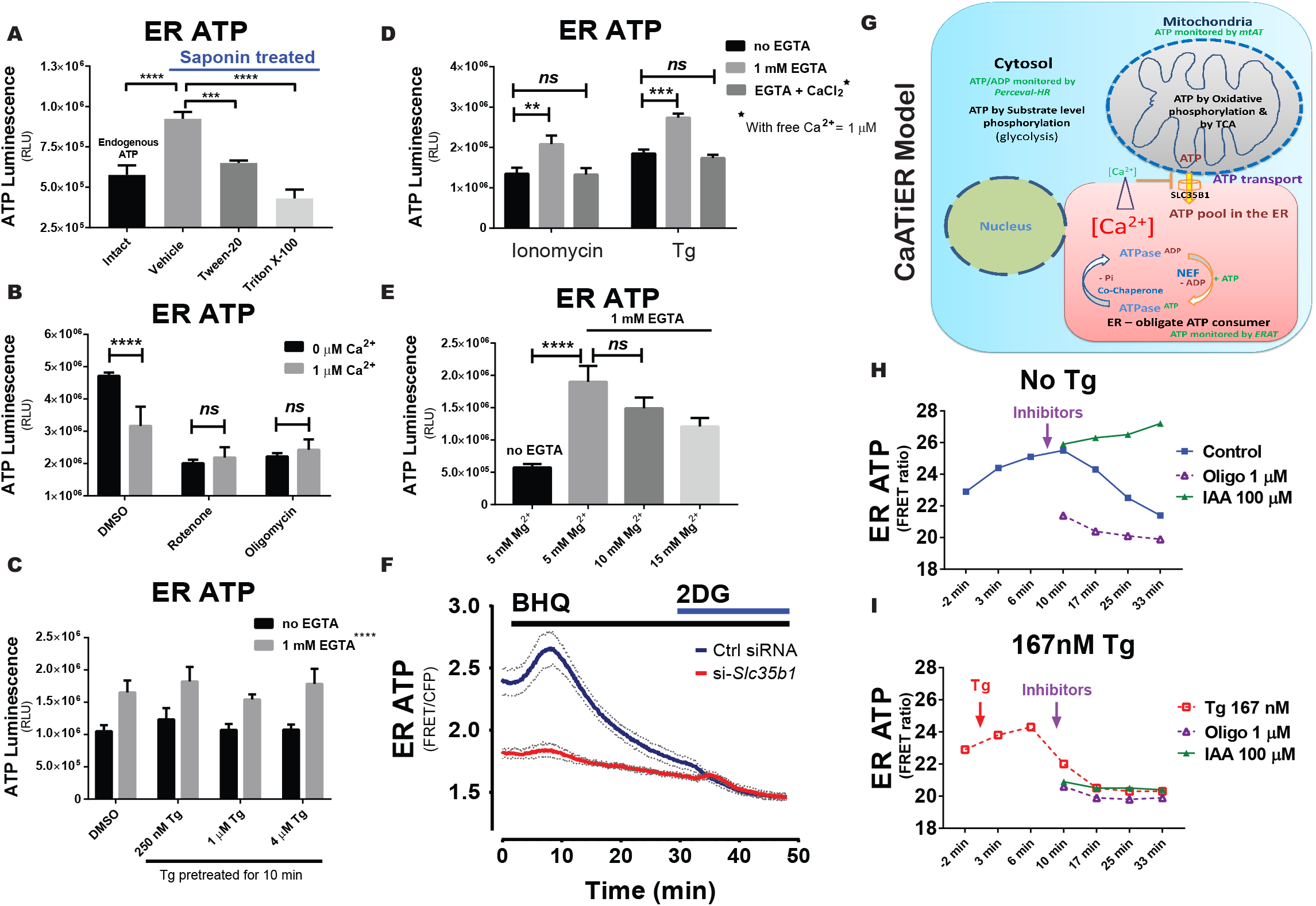
ATP transport from mitochondria to ER is inhibited by cytosolic Ca^2+^. ATP stores in plasma membrane permeabilized H9 CHO cells were measured by a reporter-free method. H9 CHO cells were permeabilized with 75 μg/mL saponin in a bathing solution (referred to as “respiration solution” hereafter). ATP production through OxPhos was supported by added pyruvate and malate (10 mM and 1 mM, respectively) as TCA substrates together with exogenous ADP (2 mM). The system was validated in two ways: First, the ATP store in the ER generated in permeabilized CHO cells was higher than that for the endogenous ATP pool detected in intact H9 CHOs (p< 0.0001 between the “*Intact*” and “*Vehicle*” group). In addition, the ER ATP store was released within 5 min after treatment with Tween-20 (0.02% vol/vol, p= 0.0007 compared to “*Vehicle*” group) or Triton X-100 (0.02% vol/vol, p< 0.0001), both after 25 min of respiration (**A**). Second, inhibition of OxPhos by oligomycin (1 μM) or rotenone (1 μM) reduces the ER ATP store to baseline levels. In addition, Ca^2+^ supplemented (in the form of CaCl_2_) at 1 μM final significantly reduced the ER ATP store (**B**). **C**. SERCA ATPase activity is blocked by pre-incubating intact H9 CHO cells with increasing concentrations of Tg (250 nM, 1 μM, and 4 μM) for 10 min. Subsequently, cells were permeabilized as described above to test if Ca^2+^ depletion affects ER ATP generated through OxPhos in permeabilized CHO cells. EGTA supplemented at 1 mM significantly increased the ER ATP store (p< 0.0001). No significant difference was observed for Tg pre-treated groups. **D**. When ionomycin (10 μM) or Tg (1 μM) were added at the same time with PM permeabilization, EGTA significantly increased ER ATP while further Ca^2+^ add-back in the form of CaCl_2_ abolished the EGTA effect on increasing the ER ATP store. **E**. Since the respiration buffer contains 5 mM MgCl2 as a basal ingredient, additional 5 to 10 mM MgCl2 was added to final concentrations of “*10 mM*” and “*15 mM*”as indicated, to test the Mg^2+^ effect on ER ATP. The difference between “*5 mM*” and “*10 mM*” was not statistically significant (p= 0.21), while EGTA supplemented at 1 mM had a significant effect (p< 0.0001) compared to 5 mM Mg^2+^ with “no EGTA”. **F**. The Ca^2+^ responsiveness of ER ATP levels by SERCA inhibition was attenuated by siRNA-mediated *Slc35b1* knockdown. HeLa cells expressing the ERAT4 reporter were treated with siRNA against *Slc35b1* for 48 hrs before cell imaging, with si-Slc35b1/siRNA control = 9.5 ± 1.4% quantified by qRT-PCR (mean ± S.E.M., n=3 samples/group). Basal ER ATP levels significantly decreased in the *si-Slc35b1* group (p< 0.001 by two-tailed t-test), compared to scrambled siRNA treated HeLa cells. Meanwhile the ER ATP decrease after BHQ addition was attenuated in the *si-Slc35b1* group. Glucose (10 mM) replacement by 2-DG (10 mM) in the end ensured cellular ATP depletion as another control. No difference was found for the two groups after 2-DG addition. **G**. The cartoon depicts the *CaATiER* model showing mitochondrial ATP transport into the ER is antagonized by Ca^2+^ on the cytosolic side of the ER membrane. **H & I**. IAA and Oligo (100 μM and 1 μM, respectively) had the characteristic effect on increasing and decreasing the ER ATP levels, respectively, in intact H9 cells (panel **H**), as shown in Figure 1. However, with Tg pre-treatment (167 nM final concentration, panel I), neither IAA nor oligomycin had an effect on ER ATP levels.

### Cytosolic Ca^2+^ inhibits ATP import into the ER

The observations presented in **Figure 2** indicate that a Ca^2+^ gradient is required to facilitate ATP transport into the ER lumen, reminiscent of the ATP/ADP exchange process mediated by the mitochondrial adenosine nucleotide transporters that is facilitated by a proton gradient across the mitochondrial inner membrane. However, the ER membrane is known to be quite “leaky” to ions, with the Ca^2+^ gradient requiring maintenance by vigorous SERCA Ca^2+^ pumping into the ER lumen. As a consequence, it is theoretically difficult for the ER membrane to build a significant chemical potential to drive ATP/ADP exchange. Alternatively, as Ca^2+^ depletion from ER lumen is inevitably coupled with a cytosolic Ca^2+^ increase in an intact cell (24), the ER ATP transporter could respond to and be regulated by cytosolic Ca^2+^, the process which we described as a *CaATiER* mechanism.

To test whether ER ATP is sensitive to cytosolic Ca^2+^, we employed an independent classical ER ATP measurement for adherent H9 cells, facilitated by a standard luminescence-based ATP assay (13, 25). The plasma membrane (**PM**) was first permeabilized with low concentrations of saponin, a cholesterol solubilizing detergent, leaving the organelle membranes intact (as these membranes contains very low levels of cholesterol) to analyze ATP transport between mitochondria and ER. ATP was generated *de novo* by supplying mitochondrial respiration substrates (pyruvate plus malate) together with 2 mM ADP in the respiration buffer. At the end of the incubation, the respiration buffer containing ATP generated by OxPhos was removed by aspiration (which accounted for > 95% of the total ATP, see below). Since the ATP pool trapped in the mitochondrial matrix cannot be eliminated technically, it is likely the mitochondrial pool of ATP contributes to the background ATP. However, we found that neither Ca^2+^ nor Tg affected mitochondrial respiration (**Sppl. Fig. S10A** to **D**). The same results further indicate that mitochondrial ATP does not change in response to Ca^2+^ or Tg used at our concentrations, as the ATP synthase is extremely sensitive to the ATP-to-ADP ratio in the mitochondrial matrix, which is further demonstrated by inhibiting respiration with carboxyatractyloside (**CAt**), an inhibitor of ANTs (**Sppl. Fig. S10A** to **D**). Theoretically, it is unlikely that OxPhos would respond to exogenous Ca^2+^ added in our assays, since greater than 1 μM Ca^2+^ is needed to open the MICU-regulated MCU gating of Ca^2+^ entry(26). Furthermore, the exogenously added ADP (2 mM) should effectively drive OxPhos-generated ATP out of the mitochondrial matrix into the ER lumen or the respiration buffer. Lastly, we verified that Ca^2+^ addition did not affect total ATP generated in the supernatant (**Sppl. Fig. S10E**) calibrated by an ATP standard curve shown alongside for the absolute amount of ATP generated in our permeabilized CHO cell system (**Sppl. Fig. S10F**).

Using this system, we confirmed that only ATP trapped in the ER compartment accounts for the changes of luminescence signals above background (i.e., *ER ATP*), which was readily liberated by brief treatment with lipid detergents (**Fig. 3A**, p< 0.001 and < 0.0001 for Tween-20 and Triton X-100, respectively). In addition, ATP generated by OxPhos from the permeabilized cells was evidenced by the higher *ER ATP* contents trapped inside PM-permeabilized cells, compared to the endogenous ATP found inside PM-intact cells (**Fig. 3A**, under “*Intact*” group). We also verified that ATP produced in this system was through OxPhos, as addition of OxPhos inhibitors at the beginning of incubation reduced ATP luminescence to background levels (**Fig. 3B**, 1 μM “*Rotenone*” or “*Oligomycin”*). Interestingly, addition of Ca^2+^ (1 μM) significantly decreased *ER ATP* (**Fig. 3B**, p <0.0001 for “*0 μM Ca^2+^” vs. “1 μM Ca^2+^” bars in the “DMSO*” group), suggesting that cytosolic Ca^2+^ inhibits ATP import from mitochondria into the ER. Alternatively, an increase in SERCA ATPase activity by increased cytosolic Ca^2+^ may reduce ATP available for the ER ATP transporter. To rule out this possibility, Tg was applied to intact H9 cells briefly before PM-permeabilization, at increasing doses from 0.25 μM to 4 μM, and the Ca^2+^ chelator EGTA was added to the respiration buffer to remove Ca^2+^ leaked from the ER into the cytosol. While 1 mM EGTA significantly increased the *ER ATP* content, addition of Tg did not have any effect (**Fig. 3C**). We also tested another Ca^2+^ releasing chemical, ionomycin – a mobile Ca^2+^ ionophore that permits Ca^2+^ transfer down the gradient, with 1 μM Tg added as a control group without pretreatment of intact cells. Again, the only effect we detected was with Ca^2+^ manipulation on the “cytosolic” side of the ER membrane, i.e. EGTA chelation increased *ER ATP* and adding CaCl_2_ back decreased the *ER ATP* (**Fig. 3D**, with free Ca^2+^ calculated to be 1 μM in presence of 1 mM EGTA). The effect of adding CaCl_2_ back was not due to increased Cl^-^ concentration, as Cl^-^ in the form of MgCl2 in great excess over CaCl_2_ only had a minor effect (**Fig. 3E**). The *ER ATP* decrease with Mg^2+^ was negligible within a physiological range from “5 *mM*” to “*10 mM*” (p =0.21 by post-doc t-test), confirming the decrease in *ER ATP* was specific to the Ca^2+^ cation. Based on a recent report identifying SLC35B1 as the putative mammalian ER ATP transporter, i.e. AXER(4), we observed that the ER ATP decrease upon BHQ treatment was attenuated when the *Slc35b1* transcript was knocked down by siRNA to 9.5%, compared to control siRNA treated cells (**Fig. 3F**). Notably, we obtained similar results as reported by *Klein et al*, despite the fact that SERCA inhibition by BHQ alone was sufficient to significantly reduce ER ATP. In addition, we observed a transient increase in ER ATP shortly (0 – 10 min) after SERCA inhibition, probably due to ER ATP usage arrest as previously reported by *Vishnu et al*. (11).

In summary, based on the above results, we propose that Ca^2+^ inhibits ER ATP uptake (**Fig. 3G**). In this *CaATiER* model, ATP from mitochondria is preferentially transported into the ER, possibly through a family of ER ATP transporter proteins represented by SLC35B1 (4), and the cytosolic Ca^2+^ concentration within this micro-domain attenuates the ATP transport mechanism. From this model, we predict that when ER ATP transport is blocked artificially by increasing cytosolic Ca^2+^, the pharmacological inhibitors for cellular ATP regeneration will lose their effects. This conclusion was successfully confirmed in intact H9 CHO cells, as treatment with low dose Tg (167 nM for ~10 min) successfully blocked IAA and oligomycin effects on ER ATP (**Fig. 3 H** versus **I**). The raw FRET signals alongside with calibrating CFP and GFP signal intensities for these treatment are shown for demonstration purpose (**Sppl. Fig. S11, A** to **D**).

### Protein misfolding in the ER increases both ER ATP dependence on OxPhos and Tg-triggered Ca^2+^ mobilization into mitochondria

Protein misfolding in the ER is a cellular stress condition characterized by activation of the unfolded protein response (**UPR**) (27). Although ER stress is typically induced by pharmacological means, e.g. with Tg to deplete ER Ca^2+^ store, or with tunicamycin (**Tm**) to inhibit N-linked glycosylation, we applied an ER stress induction model by expressing a misfolded protein in the ER lumen. ER stress was successfully induced by treating H9 CHO cells with HDAC inhibitors, such as SAHA or sodium butyrate (**NaB**), to increase F8 transcription from an HDAC-sensitive artificial promoter (14) (**Fig. 4A**). To better understand this proteostasis stress, we investigated if ATP supply and Ca^2+^ homeostasis were affected upon induction of F8. As ER stress is often associated with Ca^2+^ leak from the ER (in addition to those deliberately induced with Tg), we first validated this observation by investigating ER-initiated mitochondrial Ca^2+^ influx, by employing the mitochondrial matrix-localized Ca^2+^ reporter, mtGEM-GECO1 (28) (**Fig. 4B**). Pretreatment with 2-aminoethoxydiphenyl borate (**APB**) successfully blocked Tg-triggered mitochondrial Ca^2+^ influx, although we cannot rule the ability of APB to block other Ca^2+^ channels (29). As expected, the reporter promptly detected ionomycin-induced Ca^2+^ entry into the mitochondrial matrix (**Fig. 4C**). Although ER stress induced by SAHA treatment did not alter mitochondrial matrix Ca^2+^ levels under resting conditions, Tg-triggered mitochondrial Ca^2+^ influx was significantly more prominent in ER stressed H9 cells (**Fig. 4C** and **D**). From these results, we propose that ER protein misfolding stimulates ER to mitochondria Ca^2+^ transfer. The increased Ca^2+^ transfer into mitochondria was not due to an increased ER Ca^2+^ store since neither the O-cresolphthalein (**OCPC**) chromogenic method nor the ER Ca^2+^ reporter (*GEM-CEPIA1er*) revealed a significant increase in the ER Ca^2+^ content in SAHA-induced H9 CHO cells (**Fig. 4 E** and **F**), suggesting that the increased Ca^2+^ entry into the mitochondrial matrix under ER stress conditions is due to enhanced Ca^2+^ mobilization from another Ca^2+^ store yet to be identified (30, 31), or the close proximity between the ER and mitochondria under prolonged ER stress (32). At the same time, the oxygen consumption (by OCR) was not significantly different between ER stressed H9 CHO cells induced to express F8 and their uninduced counterparts (**Fig. 4G**). Finally, when F8 expression was induced by SAHA treatment, ER ATP homeostasis became more dependent on OxPhos, as oligomycin treatment caused greater reduction in ER ATP for SAHA-treated H9 CHO cells compared to the un-induced H9 CHO cells (**Fig. 4H**). This observation was not due to altered mitochondrial ATP supply (**Fig. 4G**) as mitochondrial ATP levels were not affected upon induction of F8 expression (**Sppl. Fig. S12A** and **B**). Lastly, we verified the oligomycin-induced decrease in ER ATP in adherent H9 CHO cells using FRET-based fluorescence microscopy (**Fig. 4 I& J**). Consistent with the ER ATP status measured by flow cytometry, the ER ATP decrease in response to complete OxPhos inhibition was significantly more pronounced in SAHA-induced ER stressed H9 CHO cells than in the vehicle treated CHO cell control group (quantified by maximal ER ATP reduction, Fig. 4K, p< 0.05). Since ATP dependence on OxPhos suggested that ER stress could have caused an “anti-Warburg effect” by inhibiting ATP production from glycolysis, we further tested if the cytosolic bioenergetic dependence on OxPhos was similarly affected by ER stress induced by Tm. Similar to the ER compartment, the cytosolic ATP/ADP ratio decreased more quickly upon OxPhos inhibition (**Sppl. Fig. S12C** and **D**), while no obvious difference was detected for the mitochondrial matrix ATP pool. To summarize, ER stress induced by a misfolded ER luminal protein causes metabolic changes, characterized by increased Ca^2+^ exchange between ER and mitochondria(24, 31) and by increased dependence on ATP produced by OxPhos.

**Figure 4.**
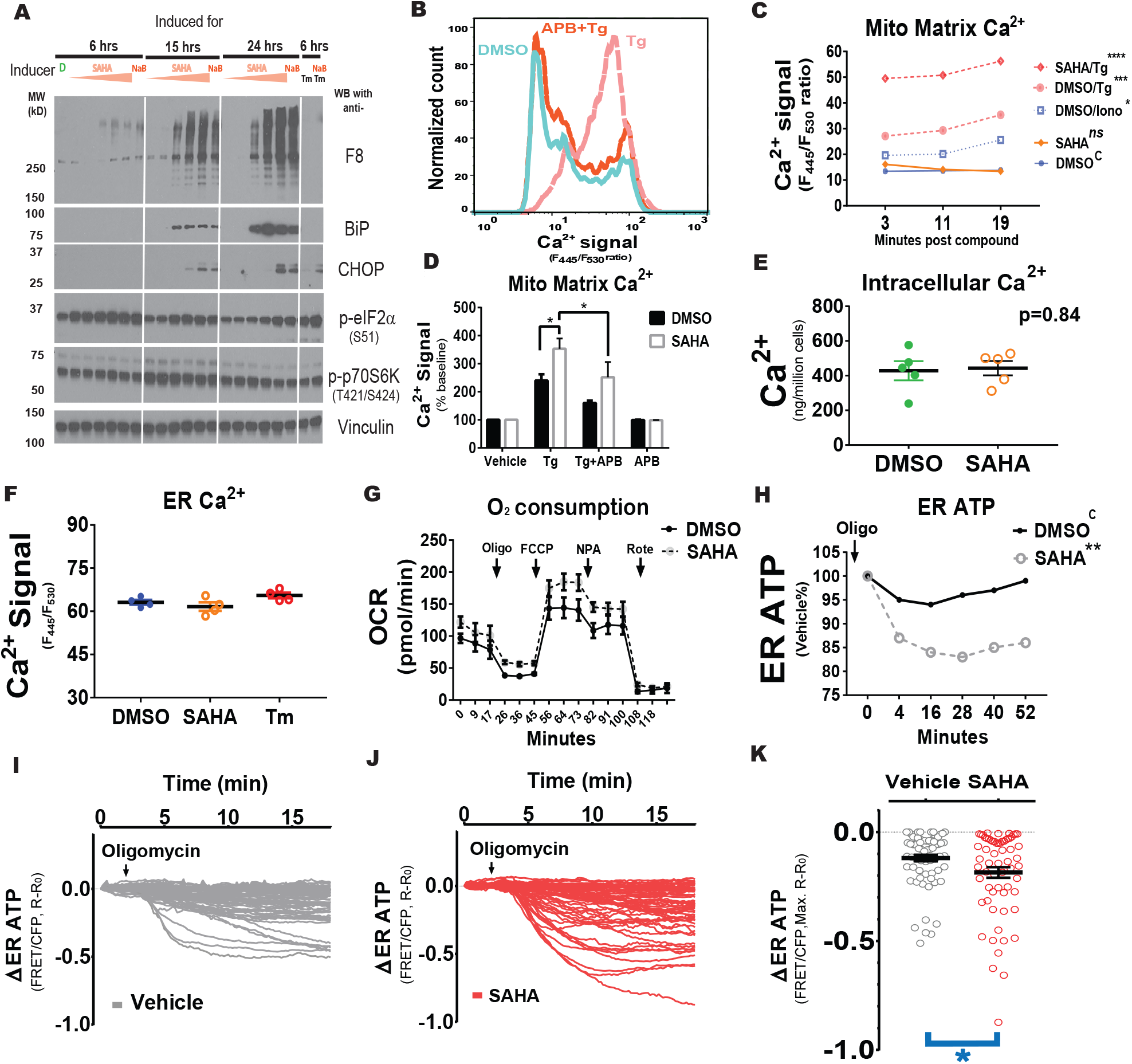
Protein misfolding in the ER increases ER ATP dependence on OxPhos and increases ER to mitochondrial Ca^2+^ trafficking. **A.** When treated with HDAC inhibitors, such as SAHA or sodium butyrate (**NaB**), F8 transcription is induced in H9 CHO cells accompanied by UPR activation (14, 15). SAHA was tested at increasing concentrations, at 50 nM, 200 nM, 1 μM, 5 μM and 20 μM, as indicated by pink arrow heads. NaB was used at 5 mM. “*D*” indicates the same volume of DMSO as a vehicle control. The last two lanes received 10 μg/mL tunicamycin (**Tm**) as positive controls for ER stress. Treatment times are labeled on the top of the Western blot panels. **B**. Representative histograms of mitochondrial Ca^2+^ influx, reported by the *mtGEM-GECO1* Ca^2+^ probe, were overlaid for visual comparison. For the particular “*Tg*” trace, H9 CHO cells were treated with 5 μM Tg for 30 min; and for the “*APB+Tg*” trace, the same cells received 50 μM of APB before Tg stimulation. **C**. While the mitochondrial Ca^2+^ concentrations show no difference at the basal level, Tg (5 μM)-triggered a higher mitochondrial Ca^2+^ spike in H9 cells with induced F8 expression. An independent group of un-induced H9 CHO cells received ionomycin (10 μM) as a positive control. **D**. Tg (5 μM)-triggered mitochondrial Ca^2+^ spikes in uninduced H9 and induced H9 cells for F8 expression are shown by bar graphs, with average signal strengths represented as percent of baseline (without Tg treatment) from four consecutive readings within 45 min. The Ca^2+^ signal from the mitochondrial matrix in ER stressed H9 cells was significantly increased compared to its vehicle-treated control group (*“SAHA*”versus “*DMSO”*, p< 0.05), both of which were partially blocked by 40 μM APB. **E**. Intracellular Ca^2+^ content was not significantly changed in H9 cells with F8 expression induced by SAHA, compared to control H9 cells as measured by the OCPC chromogenic method. **F**. ER Ca^2+^ levels in H9 CHO cells were monitored by the *GEM-CEPIA1er* probe through flow cytometry-based ratiometric analysis. SAHA (5 μM for 18 hrs) induces ER stress by increasing F8 expression in the ER, and Tm (100 ng/mL for 18 hrs) induces ER stress by blocking N-linked glycosylation on proteins translocated into the ER. ER Ca^2+^ levels in H9 CHO cells remained unchanged compared to the DMSO-treated control cells. **G**. In H9 CHO cells with induced F8 expression, the OCR was slightly increased but not significantly different, as measured by the Seahorse flux assay (n= 6 wells/group). **H**. Blockade of ATP production through OxPhos by oligomycin depletes ATP more quickly in induced H9 cells compared to un-induced H9 CHO cells, when standardized to DMSO-treated cells. **I. & J**. Traces for ER ATP in individual H9 CHO cells are shown in response to oligomycin administration (10 μM, indicated by arrow heads). In panel I, cells were treated 18 hrs with DMSO, a vehicle control for SAHA; In panel J, cells were treated 18 hrs with 5 μM SAHA. **K**. Similar to measurements made by flow-cytometry (panel **H**), when minimal ER ATP levels were compared in adherent H9 CHO cells, oligomycin caused significantly greater maximal ER ATP reduction (*, p< 0.05 by two-tailed *Student’s* t-test) in SAHA-induced H9 cells compared to “ *Vehicle*” treated H9 CHO cells.

## Discussion

ATP drives most metabolic reactions inside the cell, and its transport mechanism between mitochondria and cytosol is well characterized. In contrast, little is known about how ATP is transported into the ER, an obligate ATP consuming organelle with no known ATP regeneration machinery (5, 9).

### Mitochondria supply ATP to the ER and cellular OxPhos is enhanced by ER ATP usage

By using a series of ATP and ATP/ADP ratio reporters to monitor real-time ATP changes in sub-cellular compartments and by complementary direct measurements of ER ATP, we observed that ATP in the ER is mainly supplied from mitochondria and that ATP homeostasis in the ER is maintained through a cytosolic Ca^2+^-antagonized ATP transport mechanism on the ER membrane (**Fig. 3G**). It was surprising that the ATP level in the ER is insensitive to transient glucose deprivation and inhibition of glycolysis, which causes an immediate reduction in cytosolic ATP. This result is consistent with our conclusion that OxPhos supplies ATP to the ER and a highly efficient ATP transport mechanism exists from mitochondria to ER. Considering the close proximity between the ER and mitochondria, it is possible that ATP exiting mitochondria immediately traffics through the ER ATP transporter(s), whereas ATP produced by glycolysis is spatially more distant to be efficiently captured by the ER. In addition, supporting the symbiotic evolutionary hypothesis of mitochondria, our findings suggest that the ER takes special advantage of ATP-producing mitochondria to fulfill the energetic demands for protein folding, trafficking and secretion, reflected by the exquisite sensitivity of the ER ATP levels to OxPhos inhibition. Therefore, an intimate cross-talk between ER protein misfolding and mitochondrial function could have evolved as a bioenergetic advantage for eukaryotes to develop into multi-cellular organisms, in which cell-to-cell communications is necessitated by paracrine and endocrine signals. Based on our findings, we propose that ATP usage in the ER is a key signaling event that controls mitochondrial bioenergetics, in addition to the signaling mediated by Ca^2+^ and reactive oxygen species (**ROS**) (31).

### Cytosolic Ca^2+^ regulates ATP import into the ER

One of our most important observations is that a Ca^2+^ gradient across the ER membrane is necessary for ATP transport into the ER. Specifically, to demonstrate that cytosolic Ca^2+^ inhibits ATP import, we applied an artificial PM-permeabilized cell system, by forcing mitochondrial respiration to supply ATP, for the reason that 1) the Ca^2+^ concentration can be conveniently manipulated in this semi-*in vivo* experimental system; 2) the ATP supply is no longer limited by ADP availability which is considered a rate-limiting factor *in vivo*. Our *CaATiER* model was further validated by knocking down ER ATP transporter expression, encoded by the *Slc35b1* gene (4). Notably, the SERCA inhibitor, BHQ, induced a phenomenal ER ATP decrease which was attenuated in the *Slc35b1* knock-down HeLa cells (**Fig. 3F**). Finally, our findings demonstrate that ER Ca^2+^ release by another SERCA inhibitor (Tg) inhibited ATP transport into the ER (**Fig. 3H *versus* 3I**). The results confirm that 1) ATP is not freely permeable to the ER membrane, and 2) The ATP transport mechanism on the ER membrane (through AXER/SLC35B1) is exquisitely sensitive to the cytosolic Ca^2+^ concentration. Our findings lead us to propose that cytosolic Ca^2+^ inhibits ATP import into the ER, indicating that a Ca^2+^ responsive element faces the cytosolic surface (**Fig. 3G**), analogous to the mitochondrial Ca^2+^ uniporter (**MCU**) gating system (26, 33).

Physiological processes require exquisitely meticulous and tight regulation of ATP, i.e., cytosolic ATP abundance is sensed and converted into a Ca^2+^ chemical gradient by the coupling function of SERCA where excess cytosolic ATP will only be delivered to the ER when cytosolic Ca^2+^ is in a physiological range. In summary, we propose that cytosolic ATP usage to resolve cytosolic Ca^2+^ stress has a priority over the ATP supply into the ER. Evolutionarily, the risk of ER stress resulting from a transient ATP shortage is accommodated by bountiful ER chaperones, such as GRP78/BiP and GRP94, for which peptide binding activities are regulated by ATP (5, 7, 8), and Ca^2+^ responsive chaperones (24), such as calnexin and calreticulin (34). Previous observations associating ER Ca^2+^ release with a transient ER ATP increase (11) (and repeated in **Fig. 3F**) could reflect a temporary reduction in ER ATP consumption, rather than reflecting the luminal Ca^2+^ sensitivity of the ER ATP import machinery.

### ER protein misfolding increases the ER bioenergetic requirement for ATP from OxPhos

It has been long postulated that Ca^2+^ micro-domains formed by ER-mitochondrial contacts could potentially stimulate mitochondrial respiration to encourage efficient ATP production, especially under conditions of ER stress (31, 32). The structural proximity is maintained by the ER-mitochondria tethering molecules, such as Mitofusin 2(35, 36), while the functional cooperation through adenosine nucleotide exchange between the ER and mitochondria has not been directly proven. Uniquely here, we demonstrate that the ER preferentially depends on ATP produced from OxPhos, which was further confirmed by our observation that ER stress in H9 CHO cells causes an **“*anti-Warburg*”** effect. Whether this observation can be extrapolated to interpret cancer cell metabolism warrants further investigation. While confirming the intimate functional cooperation of ER and mitochondria through exchange of adenosine nucleotides, our findings demonstrate that increased Ca^2+^ on the cytosolic surface of the ER membrane inhibits ER ATP import and predict that the Ca^2+^ exchange domain is not compatible with the ATP exchange domain between mitochondria and the ER, suggesting the existence of either A) Two categories of contact domains between the organelles, or B) A domain that operates in two distinct modes (Ca^2+^ exchange mode versus ATP exchange mode), which are temporally separated.

ER stress caused by a protein folding overload and the subsequent accumulation of misfolded protein causes cell death under pathological conditions. ER stress-induced cell death is associated with increased oxidative stress (15, 37). OxPhos up-regulation as a feedback mechanism to increased ER ATP consumption could contribute to mitochondrial ROS generation, although altered mitochondrial function was not detected in our study (**Fig. 4G** and **Sppl. Fig. S12**). Conversely, dysfunctional mitochondria can elevate ROS production by the ER (38). In our CHO cell model system, when Tg was applied to release ER Ca^2+^, the Ca^2+^ influx into the mitochondrial matrix was significantly elevated in ER-stressed CHO cells, which could reflect the closer proximity between ER and mitochondria to ensure sufficient ATP uptake into the ER (31). *In vivo*, a similar process of IP_3_R-mediated Ca^2+^ release in response to IP3 generated by repeated plasma membrane receptor engagement may exacerbate ROS production as a result of mitochondrial Ca^2+^ overload (39).

To conclude, our results expand our knowledge of cellular bioenergetics by demonstrating that a highly efficient ATP supply mechanism exists between mitochondria and ER that is antagonized by cytosolic Ca^2+^. Our findings should stimulate research to identify and to elucidate the Ca^2+^-responsive molecular mechanism associated with the ER ATP transporter molecule(s).

## Supplemental Information

Supplemental information includes two tables and fourteen figures can be found in the attached pdf file.

## Author Contributions

Study concept and design, J.Y., A.M., R.M. and R.J.K.;

Acquisition of Data, J.Y. H.B. and M.S.;

Analysis and interpretation of data, J.Y., A.M., R.M. and R.J.K.;

Statistical analysis, J.Y. and H.B;

Technical support, M.S.;

Drafting of the manuscript, J.Y., H.B., R.M. and R.J.K.

## Acknowledgments

We thank Drs. Markus Waldeck-Weiermair and Wolfgang F. Graier from the Medical University of Graz (Austria) for sharing their critical comments and discussion of studies. We also thank Dr. Hiromi Imamura for sharing the *mtAT1.03* vector and Dr. Yi Yang for providing the *cpYFP* vector. Administrative support was kindly provided by Ms. Taryn Goode. Mrs. Yoav Altman (Flow Cytometry Core) and Leslie Boyd (Cell Imaging Core) at the SBP Medical Discovery Institute provided technical guidance and helpful discussion on imaging methodologies. Both cores receive support from the SBP NCI Cancer Center Support Grant P30 CA030199. This work was supported by the FWF Austrian Science Fund: P28529-B27 to R.M. and by NIH grants R01HL052173, R37DK042394, R24DK110973, R01DK103185, R01DK113171 and R01CA198103 to R.J.K.

