## Supplemental information for "Mitochondria supply ATP to the ER through a mechanism antagonized by cytosolic Ca^2+^"

Running Title: Regulated ATP transport into the ER lumen from mitochondria.

Keywords: ER Stress, protein folding, nucleotide transport, bioenergetics, ATP.

### Supplemental Information

#### Methods details

##### Construction and validation of H9 CHO cell line

The F8 inducible H9 CHO cell system was described previously (14, 15). Expression of F8 was confirmed by Western blot using specific monoclonal antibodies – anti-F8 (Green Mountain Antibodies, GMA 012, murine IgG1), anti-BiP (Cell Signaling Technology, CST 2177, rabbit IgG), anti-CHOP (Santa Cruz Biotechnology, SC 575, rabbit IgG), anti-phospho-eIF2α at Ser51 (Cell Signaling Technology, CST 3597S, rabbit IgG), anti-phospho-p70S6K at Thr421/Ser424 (Cell Signaling Technology, CST 9204, rabbit IgG), and anti-VINCULIN (Sigma, V 9131, murine IgG1).

##### Construction of novel reporter cell lines

CHO cells were transfected with the reporter plasmids using the Amaxa Nucleofector II electroporation system in the Mirus Ingenio solution (Cat # MIR 20114). For reporter plasmids with G418 resistance, a brief G418 selection (0.5 mg/mL) was applied on transfected CHO cells for one week. The population of surviving cells was allowed to recover for two weeks before they were used for ER ATP analysis by flow cytometry. For reporter plasmids without a mammalian selection marker, a puromycin resistance cassette (~1.23 kb, amplified from plasmid pGL4.21) was co-transfected using the *Amaxa Nucleofector II* electroporation system in the *Mirus Ingenio* solution. After a brief puromycin selection (5 µg/mL for five days), cells were allowed to recover for two weeks before they were used for fluorescence ratio analysis by flow cytometry.

More specifically, plasmids encoding the *ER-RFP* reporter (ER-mRFP, #62236), the *PercevalHR* reporter (*GW1-PercevalHR,* #49082), the *CEPIA1er* reporter (p*CIS GEM-CEPIA1er,* #58217) and the *mtGEM-GECO1* reporter (*CMV-mito-GEM-GECO1*, #32461) were purchased from *Addgene*. The plasmid encoding the *ERAT4.01*$^{N7Q}$ reporter was purchased from *Next Generation Fluorescence Imaging* Company (NGFI, Austria). The plasmid encoding the *TagRFP* reporter was purchased from *Evrogen* (pTagRFP-C, #FP141). The plasmid encoding the *mtAT1.03* reporter was a kind gift from Dr. Hiromi Imamura at Kyoto University. The plasmid encoding the *cpYFP* reporter was a kind gift from Dr. Yi Yang at the East China University of Science and Technology, and the plasmid encoding the *D1ER* reporter was a kind gift from Dr. Demaurex at Universitè de Genève, Switzerland. Refer to **Supplementary Table I** for a summary of the reporter proteins' subcellular localization and their reporting specificity.

##### Confocal microscopy

For live cell imaging of subcellular localization of ERAT in H9 CHO, targeting to the ER was validated using an array confocal laser scanning microscope (ACLSM) built on a fully automated inverse microscope (Axio Observer.Z1, Zeiss, Göttingen, Germany), using a 100x objective (Plan-Fluor 100 x/1.45 oil, Zeiss). CHO cells expressing ERAT were additionally transfected with ER-RFP (40), and either treated with vehicle (DMSO) or 5 µM SAHA for 18 hrs. Live cells were excited using diode lasers (Visitron Systems, Puchheim, Germany): CFP of ERAT was excited at 445 nm (50 mW), RFP of ER-RFP was excited at 561 nm (50 mW). Emissions were collected using the emission filters ET480/40 for CFP and E570LPv2 for RFP, respectively (Chroma Technologies Corporation, VT, USA). All images were captured at a binning of 2 using a photometrics CCD camera (CoolSnap HQ2, Photometrics, Arizona, USA), and images were processed using ImageJ software.

For fixed H9 CHO cells after immunostaining with anti-PDIA6 antibody, confocal images of stably transfected CHO cells were taken on an Olympus Fluoview 1000 platform, using the following excitation and emission filters - for DAPI, Ex 405 nm/Em 461 nm; for ERAT protein, Ex 488 nm/Em 519 nm; for PDIA6 (immuno-detected by Alexa-594 conjugated secondary antibody), Ex 561 nm/Em 618 nm. Objective lenses of 60x (PlanApo N, N.A = 1.42 oil) and 100x (UPlanSApo N.A. = 1.40 oil) were used.

**Quantification of ER ATP and $Ca^{2+}$ levels by fluorescence ratiometry analysis in live cells**

Analyses of ER ATP levels in INS-1 832/13 (INS-1) cells were facilitated with the ERAT 4.01 reporter introduced by conventional lipofectamine-transfection. Imaging of ER $Ca^{2+}$ levels was performed using the ER targeted $Ca^{2+}$ reporter D1ER. Prior to measurements, cells were equilibrated for 30 min using storage buffer composed of (mM): 138 NaCl, 5 KCl, 2 $CaCl_2$, 1 $MgCl_2$, 10 HEPES, 2.6 $NaHCO_3$, 0.44 $KH_2PO_4$, 0.34 $Na_2HPO_4$, 10 D-glucose, 2 L-glutamine, with the following supplements (vol/vol): 0.1% vitamins, 0.2% essential amino acids, and 1% penicillin–streptomycin, pH adjusted to 7.4 with NaOH. The experimental buffer used for the perfusion of the cells during the fluorescence microscopic experiments was composed of (mM) 138 NaCl, 5 KCl, 1 $MgCl_2$, 10 HEPES, 10 glucose, either with 2 $CaCl_2$ or 0.1 EGTA with no added $CaCl_2$, pH adjusted to 7.4 with NaOH. During the experiment, buffers were exchanged using a flow chamber, connected to a gravity-based perfusion system (NGFI, Graz, Austria) and a vacuum pump (Chemistry diaphragm pump ME 1c, Vacuubrand, Wertheim, Germany).

CHO cells stably expressing the ERAT4.01$^{N7Q}$ reporter were directly analyzed after removing the alpha MEM medium. Cells were cultured and analyzed in µ-slide 4 wells (ibidi GmbH, Planegg, Germany). For the experiments, alpha MEM was removed and replaced by experimental buffer composed of (mM): 138 NaCl, 5 KCl, 1 $MgCl_2$, 10 HEPES, 10 glucose, 2 $CaCl_2$, 2 pyruvate and 4 glutamate, pH adjusted to 7.4 using NaOH.

All fluorescence microscopic experiments were performed using an iMic inverted and advanced fluorescent Microscope with a 40x magnification objective (alpha Plan Fluor ×40, Zeiss, Göttingen, Germany) and a motorized sample stage (TILL Photonics, Graefling, Germany). Excitation was performed at 430 nm (Polychrome V, Till Photonics), emissions were collected simultaneously at 475 nm for CFP and 525 nm for YFP, respectively, using a beam splitter. Images were captured using a binning of 2 with a CCD camera (Allied Vision Technologies, Stadtroda, Germany) and analysis was performed using Live Acquisition software (TILL Photonics). Representative ratio images were created using MetaMorph microscopy automation and image analysis software (Molecular Devices, Sunnyvale, CA, USA).

**ER and mitochondrial ATP level analysis by flow cytometry**

For FRET-based ATP determination, a Novocyte 3000 flow cytometer (ACEA BioSciences) was used to record ERAT fluorescence at channels with Ex/Em filters set as following: 1). 405 nm/ 445 (band width: 45) nm; 2). 405 nm/ 530 (band width: 30) nm. ER and mitochondrial ATP levels for individual cells were defined as the ratio of fluorescence intensity of channel 2 (Ex/Em: 405/530) divided by that of channel 1 (Ex/Em: 405/445), similar to parameters used previously (10, 11). In addition, fluorescence intensities of the following channels were recorded for probe abundance quantification, and for data validity verification: 1) 405 nm/ 572 (band width: 28) nm; 2) 488 nm/ 530 (band width: 30) nm; and 3) 488 nm/ 572 (band width: 28) nm.

**Cytosolic ATP/ADP ratio analysis by flow cytometry**

For cytosolic ATP/ADP ratio determination in H9 CHO cells expressing the *PercevalHR* reporter, an LSR Fortessa flow cytometer (BD Biosciences) or a Novocyte 3000 flow cytometer (ACEA BioSciences) were used to record fluorescence intensity of channels with Ex/Em as following: 1) 405 nm/ 525 nm (band width: 50 nm); and 2) 488 nm/ 510 nm (band width: 25 nm). The cytosolic ATP/ADP ratio for individual cells is defined as the ratio of fluorescence intensity of channel 2 (Ex/Em - 488/510) divided by that of channel 1 (Ex/Em - 405/525), similar to previously described (17). In addition, as fluorescence signals from the *PERCEVAL-HR* protein are known to be sensitive to cytosolic pH changes upon compound addition, the ratiometric $F_{525}/F_{510}$ was corrected by simultaneously measuring pH changes using the cpYFP overexpressing H9 CHO cells using a first-order correction (41).

**Mitochondria $Ca^{2+}$ influx analysis by flow cytometry**

CHO cells were transfected with the *mtGEM-GECO1* plasmid as described above, and after brief G418 selection for a week, cells were allowed to recover for two weeks before they were used for mitochondria $Ca^{2+}$ influx analysis by flow cytometry using a Novocyte 3000 flow cytometer. For mitochondrial matrix $Ca^{2+}$ level, a ratiometric measurement was derived using fluorescence intensity of channel 1 with Ex/Em of 405 nm/ 445 nm (band width: 45 nm) divided by that of channel 2 with Ex/Em of 405/530 nm (band width: 30 nm), similar to a procedure previously described (28).

**ER luminal $Ca^{2+}$ analysis by flow cytometry**
CHO cells were transfected with the *GEM-CEPIA1er* plasmid as described above, and after brief puromycin selection (5 µg/mL for five days), fifteen percent of the transfected cells were positive for the *CEPIA1er* reporter as quantified by flow cytometry analysis. Cells were allowed to recover for two weeks before they were used for ER lumen $Ca^{2+}$ analysis by flow cytometry facilitated by a Novocyte 3000 flow cytometer. For $Ca^{2+}$ determination, a ratiometric measurement was derived using fluorescence intensity of channel 1 with Ex/Em of 405 nm/ 445 nm (band width: 45 nm) divided by that of channel 2 with Ex/Em of 405/530 nm (band width: 30 nm), similar to a procedure previously described (22).

**Flow data analysis by FlowJo software**
Flow cytometry data recorded by cytometers were saved as FCS 3.1 files and analyzed by FlowJO software (Version: 10.2, FlowJo LLC) on a Mac platform. Briefly, cells were first gated by forward light scatter and side light scatter (FSC-H and SSC-H, respectively) to ensure that only fluorescence signals from live singlet cells were analyzed. Within the population, subsets of reporter-positive and reporter-negative populations were further differentiated by plotting the fluorescence intensity at designated channels. A ratio parameter was further defined using the FlowJo's "*Derived Parameters*" function, as defined above for each reporter. Normality of the derived parameter was subsequently checked by plotting a histogram for every ratiometric parameter. Furthermore, the geometric mean of fluorescence intensity (gMFI) was applied for population statistics analysis, as ratiometric parameters usually follow a log-normal distribution. Typically, statistics from a reporter positive population of more than 3000 singlet cells at any given time point were used to ensue data reproducibility.

**Effect of pharmacological reagents on ER and mitochondria ATP levels**
Mito-toxins and glycolysis inhibitors were diluted to 100x stock concentrations in DMSO or PBS. All chemical inhibitors were purchased from Sigma or Thermo Fisher unless indicated otherwise. On the day of flow analysis, 10 µL of compounds were aliquoted into Falcon polypropylene tubes (Corning Cat # 352063) before 1 mL of cell suspension was added to make the final concentration. In any particular experiment where multiple compounds were used for compound effects comparison, e.g. Fig. 1E and 1F, all compounds were pre-pipetted into the Falcon tubes before cell addition. To ensure population homogeneity, cells from the same tubes were aliquoted to the compound-containing tubes or to the vehicle-containing tubes for data collection. A graphic depiction of the standard assay procedure is shown in **Sppl Fig.S13**.
Furthermore, based on the observation that H9 cells with high ERAT4 reporter expression have improved data normality for FRET ratio (**Sppl. Fig. S14, A&C**), we generated a single H9-D2 CHO cell clone with unanimous high levels of ERAT4 expression (**Sppl. Fig. S14, B&D**), by seeding single cells into multiple 96-well cell culture plates, followed by manual selection of clones with high YFP fluorescence intensities. Data sphericity for FRET ratio of H9-D2 CHO cell was further confirmed by flow cytometry based FRET signal analysis, as described above.

**Cellular metabolic flux analysis by Seahorse XF24 platform**
Cellular oxygen consumption rate (**OCR**) and extracellular acidification rate (**ECAR**) were analyzed on a Seahorse XF24 analyzer (*Seahorse BioSciences*). H9 CHO or CHO-DUK cells were seeded onto a XF24 cell culture microplate 4-5 hrs before the assay to allow cell attachment to the plate. OCR and ECAR were measured following XF24's standard operating procedures as per manufacturer's manual,

facilitated by a XF24 extracellular flux assay kit (*Seahorse BioSciences*, Part # 100850-001). In addition, to measure OCR and ECAR in F8-induced H9 CHO cells, the subject CHO cells were first induced by 5 µM SAHA for 21 hrs in a 10-cm cell culture dish before re-plating into a XF24 cell culture microplate, to ensure equal cell input to the control un-induced H9 CHO cells.

For permeabilized CHO cell respirometry, cells were permeabilized with rPFO reagent (Agilent #102504-100) in a cytosol-like buffer containing 70 mM sucrose, 220 mM mannitol, 10 mM $KH_2PO_4$, 5 mM $MgCl_2$, 2 mM HEPES, 1 mM EGTA and 0.2% BSA(w/v), with pH adjusted to 7.4 by KOH.

The standardized procedure measuring mitochondrial respiration on an XF platform can be found at the following website: ***https://www.agilent.com/cs/library/technicaloverviews/public/5991-7157EN.pdf***

**ER ATP assay in saponin-permeabilized H9 CHO cells measured by luciferase assay**

H9 CHO cells were seeded onto 96-well plates at a density of fifty thousand cells per well and were allowed to attach and grow overnight to reach eighty to ninety percent confluency. On the day of assay, cells were first permeabilized with artificial cytosol-like buffer containing 75 µg/mL saponin. The buffer contains 70 mM sucrose, 220 mM mannitol, 10 mM $KH_2PO_4$, 5 mM $MgCl_2$, 2 mM HEPES, 1 mM EGTA (unless when it was intentionally omitted) and 0.2% BSA(w/v), with pH adjusted to 7.4 by KOH. In addition, 10 mM pyruvate and 1 mM malate plus 2 mM ADP were added to maintain mitochondria respiration upon membrane permeabilization. $Ca^{2+}$ concentrations in cytosol-like buffer were controlled by adding increasing amounts of $CaCl_2$. Free [$Ca^{2+}$] was estimated according to a protocol previously described (42) (**Supplementary Table II**). H9 cells were further incubated at 37 ºC in a cell culture incubator for 20-30 min, before the buffer was removed by taping the cell culture plate on a stack of dry paper towels. *ATPLite* kit (Perkin Elmer, Cat # 6016943) was used to quantify the ATP store in permeabilized H9 cells, following the manufacturer's protocol. Bioluminescence signals were recorded on a *SpectraMax i3x* multi-mode detection platform (Molecular Devices), with an integration time of 500 milliseconds per well. ATP measurements from eight to ten wells were averaged to plot the final graph under all conditions. Tg (0.25 - 1 µM) and oligomycin (1 – 2.5 µM) were used in inhibit SERCA and ATP synthase respectively, as indicated.

In addition, the same *ATPLite* kit was used to measure total cellular ATP. For **Fig. 1H** and **Sppl. Fig. S6A** and **S7A**, H9 CHO cells were incubated for 1 hr in serum-free DMEM medium containing substrates supporting only OxPhos (Seahorse BioSciences, Cat. #100965 with 1 mM sodium pyruvate supplemented), in experiments where glucose supplementation effect was tested.

**SLC35B1 knockdown in HeLa cells**

For siRNA mediated knockdown of SLC35B1 in HeLa cells, the following siRNA sequence, targeting the *Slc35b1* 3'-UTR, was used: 5'-GAG ACU ACC UCC ACA UCA A dTdT-3'. Control cells were treated with a scrambled siRNA of the following sequence: 5'-AGG UAG UGU AAU CGC CUU G dTdT-3'. Cells were transfected at a confluency of ~70% using TransFast transfection reagent (Promega GmbH, Mannheim, Germany) according to the following protocol: 1.5 µg ERAT4.01 plasmid, 0.12 nmol of siRNA against SLC35B1 UTR or control siRNA and 3 µl of TransFast were added to 1 ml of FCS free DMEM medium. Mixture was incubated for 15 min at room temperature before applying onto adherent HeLa cells, replacing cell growth media containing 10% FCS. After 4 hrs, transfection mixture was exchanged for DMEM plus 10% FCS. Cells were incubated for another 48 hrs in a humidified incubator at 37°C prior to imaging analysis.

**Detection of intracellular $Ca^{2+}$ levels by chromogenic assay in CHO lysates**

Total intracellular $Ca^{2+}$ levels were measured by the O-Cresolphtalein (OCPC) chromogenic method, using components provided in a $Ca^{2+}$ Assay Kit (Adipogen Corp., Cat # JAI-CCA-030). Briefly, H9 CHO cells were grown to 70% confluency in 10 cm cell culture dishes and induced to express F8 with 5 µM SAHA for 22 hrs. Cells were trypsinized, gently centrifuged and re-suspended in 1mL complete culture medium. After cell density quantification, six million cells were dispensed into a separate set of 1.5 mL Eppendorf tubes for $Ca^{2+}$ quantification. Cells were pelleted by centrifugation, washed once with $Ca^{2+}$-free HBSS, and lysed immediately in 120 µL of 3% trichloroacetic acid on ice for 30 min, with intermittent vortexing. Cell lysates were further clarified by centrifugation at 6,000 rpm for 15 min using a bench-top Eppendorf Centrifuge (Model # 5415R). Cleared supernatants were used for OCPC-based $Ca^{2+}$ level detection as per manufacturer's instructions. Optical densities (OD at 570 nm) were read and recorded by a *VERSAmax* microplate reader (Molecular Devices) and $Ca^{2+}$ concentrations were determined by the $Ca^{2+}$ standard curve provided in the OCPC kit.

**Western blot analysis**

CHO cells were grown on multiple-well plates and treated with compound in alpha MEM growth medium (14). Cells were harvested by direct lysis with RIPA buffer (150 mM NaCl, 50 mM Tris pH 7.4, 1% Triton X100, and 0.5% Sodium Deoxycholate). Cell lysates were pre-cleared by centrifugation and protein concentration were determined by Bio-Rad's Dc protein assay kit. Total cellular protein (10 – 20 µg) was loaded onto a Bio-Rad's precast SDS-PAGE gel for electrophoresis and subsequent semi-dry transfer to a nitrocellulose membrane, as previously described (15). Primary antibodies used for immunoblotting are summarized in "*Construction and validation of H9 CHO cell line*" section.

**Quantification and Statistical Analysis**

Flow cytometry data recorded by cytometers were saved as FCS 3.1 files and analyzed by FlowJo software (Version: 10.2, FlowJo LLC) on a Mac platform. A ratio parameter was defined using the FlowJo's "*Derived Parameters*" function, as defined above for each reporter. Normality of the derived parameter was subsequently checked by plotting a histogram for every ratio parameter. Furthermore, the geometric means of fluorescence intensities (gMFI) were applied for population statistics analysis, as ratiometric parameters usually follow a log-normal distribution. Typically, statistics from a reporter positive population of more than 3000 single cells at any given time point were used to ensure data reproducibility.

Colocalization by *Pearson correlation* analysis of CHO cells expressing $ERAT4.01^{N7Q}$ and *TagRFP* was performed using ImageJ software, with plugins "*Colocalization Test*" and "*Fay Randomization*" algorithms. Prior to the analysis, images were further 2D-deconvoluted using MetaMorph software (Molecular Devices, San Jose, USA).

Compartmental ATP and/or $Ca^{2+}$ levels were compared using the "*Ordinary Two-way ANOVA*" function provided by GraphPad's Prism software (Ver: 7.0), with alpha level of 0.05. Time and compounds were assumed two independent parameters for ANOVA analysis. *Dunnett's* multiple comparisons test was subsequently performed for comparison between individual groups versus diluent control (DMSO or PBS), or control group as indicated for particular experiments.

Statistical significance was expressed as following: n.s: - not significant; * - $p \leq 0.05$; **- $p \leq 0.01$; ***- $p < 0.001$; **** - $p < 0.0001$.

**Supplementary Table I. Summary of reporters used in CHO cells and their intended specificity.**

| Name | Reporter for | Compartment Localization | Imaging property | Reference |
|---|---|---|---|---|
| mtAT 1.03 | ATP level | Mito matrix | FRET | *Imamura et al.,* Ref. 10 |
| ERAT 4.01 N7Q | ATP level | ER lumen | FRET | *Vishnu et al.,* Ref. 11 |
| TagRFP | Cytosolic location | Cytosol | Fluorescent | *Merzlyak et al.*, Ref. 16 |
| Perceval HR | ATP/ADP ratio | Cytosol | Ratiometric | *Tantama et al.*, Ref. 17 |
| D1ER | $Ca^{2+}$ level | ER lumen | FRET | *Palmer et al.*, Ref. 21 |
| GEM-CEPIA1er | $Ca^{2+}$ level | ER lumen | Ratiometric | *Suzuki et al.*, Ref. 22 |
| mtGEM-GECO1 | $Ca^{2+}$ level | Mito matrix | Ratiometric | *Zhao et al*., Ref. 28 |
| ER-RFP | ER localization | ER lumen | Fluorescent | *Snapp et al.*, Ref. S40 |
| cpYFP | pH level | Cytosol | Ratiometric | *Zhao et al.*, Ref. S41 |

**Supplementary Table II. Free $Ca^{2+}$ concentration estimates for $CaCl_2$ containing respiration buffers (37ºC)**

| Other chelating ingredients | $CaCl_2$ (mM) | $Ca^{2+}$ (nM) |
|---|---|---|
| EGTA 1 mM and $Mg^{2+}$ 5 mM ATP = ~ 0.2 mM pH ~7.4 | 0.100 | 17 |
| | 0.250 | 50 |
| | 0.500 | 150 |
| | 0.750 | 453 |
| | 0.800 | 605 |
| | 0.875 | 1,000 |
| | 0.900 | 1,300 |
| | 0.950 | 11,100 |

#### Figure Legends for Supplemental Figures

##### Figure S1

**The ERAT probe is localized to the ER lumen.**

**A**. Confocal microscopy confirms ER localization of the ER ATP reporter, ERAT, in fixed H9 CHO cells. A representative confocal micrograph shows a high degree of co-localization of ERAT fluorescence in green with PDIA6 immunofluorescence in red. Nuclei were counter-stained with DAPI in blue. (Scale bar: 20 µm)

**B**. F8 expression upon SAHA treatment does not alter ERAT's co-localization with PDIA6. (Scale bar: 10 µm)

##### Figure S2

**The ERAT probe is localized to the ER lumen.**

**A.** Confocal microscopy confirms ER localization of the ER ATP reporter, ERAT, in live H9-D2 CHO cells. A representative confocal micrograph shows a high degree of co-localization of ERAT fluorescence in green with ER-RFP maker fluorescence in red. In addition, there are ERAT-positive cells without ER-RFP expression.

**B.** F8 expression upon SAHA treatment does not alter ERAT's co-localization with ER-RFP maker, in live H9-D2 CHO cells. Yellow arrowhead indicates a protein aggregate of RFP in cytosol. (Scale bar: 20 µm; the same scale bar is used for **A** and **B** panels)

##### Figure S3

**2-DG treatment does not reduce OxPhos of H9 CHO cells.**

**A.** 2-DG treatment for ~ 30 min (20 mM final concentration) has no effect on cellular oxygen usage. Oxygen consumption rates (**OCR**, in pMole/min) in H9 CHO cells were measured by an *XF-24* platform before serial injections of the OxPhos inhibitors (as shown in main Fig. 1D). OCR values from individual wells for both groups are shown here as scattered plots with Mean ± SEM. The "*n.s.*" note indicates no significance detected.

**B.** In the same metabolic flux experiment as shown in main Fig. 1D, 2-DG treatment immediately attenuated extracellular acidification rate (ECAR, in milli pH per minute), a surrogate measure designed to reflect cellular glycolysis efficiency.

#### Figure S4

**ER localization of ERAT probe is not affected by brief treatment with bio-energetic inhibitors.**

Confocal microscopy confirms ER localization of the ERAT reporter in live H9-D2 CHO cells, both before and after treatment with bio-energetic inhibitors for 30 min. A representative confocal micrograph shows distinct compartmentation of ERAT fluorescence in green, in contrast to cytosol-targeted TagRFP fluorescence in red. Bio-energetic inhibitors used here are **A**. DMSO as vehicle control (0.01%, vol/vol); **B**. 10 mM 2-DG; **C**. 3 µM FCCP; **D**. 3 µM oligomycin; **E**. 5 µM rotenone; **F**. 100 µM IAA.

(**G**). Analysis of Pearson correlation coefficient shows no change in ERAT fluorescence pattern before and after inhibitor treatment.

#### Figure S5

**The dynamic range of the ER ATP change was determined by flow cytometry.**

In an effort to determine the dynamic range of the FRET ratio in response to ER ATP change, H9 CHO cells were first treated with DMSO **A**. (1% by volume) or 2-DG **B**. (20 mM) for 2 hrs, before the cells were trypsinized and analyzed by flow cytometry. After two readings by flow (0 min and 7 min), half of the cells in suspension were further treated with 1 µM oligomycin, indicated by the "Oligo" with arrowhead. The "Vehicle" group received DMSO as a control. A raw value of ~21 was determined to be the lowest range after the complete ATP regeneration machinery was shut off.

#### Figure S6

**Glucose supplementation increases total cellular ATP while 2-DG decreases the cytosolic ATP-to-ADP ratio.**

**A.** Glucose supplementation (5 mM x1 hr) in basal medium with only OxPhos substrates increases the total cellular ATP content. Basal medium is DMEM medium with only OxPhos substrates (Seahorse BioSciences, Cat. # 100965 with 1 mM sodium pyruvate supplemented). **B.** In complete H9 CHO medium (Alpha MEM, no FBS supplemented), 2-DG (20 mM) greatly reduces total ATP content in H9 cells, while oligomycin (1 µM) slightly reduces total ATP content.

**C.** 2-DG treatment reduces cytosolic the ATP-to-ADP ratio. 2-DG (20 mM) was injected at 0 min. The ATP/ADP ratios were significantly decreased in H9 CHO cells that received 2-DG.

**D.** Oligomycin treatment (1 µM) does not significantly alter the cytosolic ATP-to-ADP ratio.

For panels A & B, total cellular ATP contents were measured by *ATPLite* kit. For panels C & D, cytosolic ATP/ADP ratios in H9 CHO cells were monitored by the *PercevalHR* probe through flow cytometry-based ratiometric analysis. The ratiometric *PercevalHR* parameter values were further corrected by the *cpYFP* ratiometric reading from cells treated simultaneously with the same toxins, in a separate set of H9 CHO cells, for pH correction.

#### Figure S7

**Glucose supplementation does not alter ER ATP levels and 3-Bromopyruvate increases ER ATP levels.**

**A.** Glucose supplementation (5 mM x 1hr) in basal medium with only OxPhos substrates has no effect on ATP levels in the ER, while oligomycin reduces ER ATP. The glucose supplemented group is represented by a green solid line, and basal media group (DMEM medium with only OxPhos substrates, Seahorse BioSciences, Cat. # 100965 with 1 mM sodium pyruvate supplemented) is represented by a blue dotted line. The oligomycin treated group is shown by a red solid line.

**B.** 3-Bromopyruvate (BrP) was injected at the indicated concentrations and ER ATP levels were monitored in H9 CHO cells expressing the ERAT reporter. ER ATP levels increased briefly after BrP injection at 20 µM and 100 µM.

**C.** ER ATP levels in H9 CHO cells in response to IAA and BrP (both at end concentration of 100 µM) were compared in the same experiment. Both compounds increased ER ATP levels detected by the FRET-based ERAT reporter.

**D.** OxPhos inhibitors (oligomycin and rotenone) reduce ER ATP in DUK-CHO cells that do not express human F8. In contrast, glycolysis inhibitors, represented by 2-DG and IAA at the indicated concentrations, increase ER ATP levels. Statistical significance values are labeled in the legend for individual compounds, relative to the “DMSO” group. Two-way ANOVA was applied for statistical analysis of geometric means of fluorescence intensity (gMFI), with significance levels expressed as: n.s: - not significant; * - $p \leq 0.05$; **- $p \leq 0.01$; ***- $p < 0.001$; **** - $p < 0.0001$.

#### Figure S8

**ER ATP levels in INS1 cells decrease after BHQ-mediated SERCA inhibition and recovery takes time.**

**A.** Experimental scheme for results shown in panels **B** & **C**. Briefly, INS-1 cells were first treated with 15 µM BHQ for 30 min to reduce ER ATP levels, and after BHQ removal, the cells were allowed to recover in medium for a period of time up to 4 hrs and their ER ATP levels were measured by ERAT FRET ratio.

**B.** Basal ERAT FRET Ratio values of INS-1 cells, either untreated or at different periods of time after BHQ-induced ER ATP depletion. There is notable ER ATP recovery in INS-1 cells starting at 30 min after BHQ removal.

**C.** Maximal decrease in ER ATP from basal FRET ratios is shown for INS-1 cells after a second stimulation with 15 µM BHQ. Note that the ER ATP levels were recorded from the same batch of INS-1 cells used in panel B, by the same color code.

**Figure S9**

**Tg and CPA decrease ER $Ca^{2+}$ and ATP in H9 CHO cells due to SERCA inhibition.**

**A.** Treatment with cyclopiazonic acid (CPA, at 50 µM) or Tg (1 µM) for 18 min decreases ER ATP levels. ER ATP was estimated by flow cytometry using the ERAT reporter.

**B.** ER $Ca^{2+}$ levels in H9 CHO cells were monitored by the GEM-CEPIA1er probe by flow cytometry-based ratiometric measurements. Tg (500 nM, injected at 0 min) depleted ER $Ca^{2+}$ within ~ 9 min after addition, a significant effect compared to DMSO-treated control cells.

**C. & D.** Upon treatment with Tg (0.25 µM, added at 0 min), the ER ATP levels in H9 CHO cells decrease significantly faster (**C**) than the mitochondrial ATP (**D**). ER ATP levels were measured by the ERAT probe and mitochondrial ATP was measured by the mtAT 1.03 probe. ATP levels in the two compartments were measured in parallel in the two sets of H9 CHO cells using the same FRET versus CFP ratio ($F_{530}/F_{445}$). The ATP levels measured by FRET ratios were further used to calculate Tg-induced ATP decrease standardized to the respective DMSO-treated control cells, and ATP levels were expressed as percentage of the "DMSO" group (% of DMSO as shown in main Fig. 2G).

**Figures S10**

**Permeabilized H9 CHO cells produce ATP through OxPhos and ATP is detected in the respiration buffer.**

**A.** $Ca^{2+}$ supplementation does not alter mitochondrial respiration of PM-permeabilized H9 CHO cells. Oxygen consumption rate (OCR in pMole/min) was measured using a Seahorse XF-24 platform. For the "$Ca^{2+}$" group (n=10 wells), wells received 1 µM $Ca^{2+}$ in the form of $CaCl_2$ injected after the fourth measurement, while the vehicle control group (n=10 wells) received buffer alone. Subsequently, both groups received sequential injections carboxyatractyloside (**CAt,** 3 µM), FCCP (1 µM), and finally rotenone (1 µM) plus antimycin A (10 µM). Data are represented as Mean ± SEM.

**B.** $Ca^{2+}$ supplementation does not alter mitochondrial respiration in PM-permeabilized H9 CHO cells. Data from panel (**A**) were used to calculate group averages after each inhibitor treatment, which are shown as bar graphs, with SEM represented by whiskers. Two-way ANOVA detected no effect of $Ca^{2+}$ at 1 µM, with a p-value of 0.61.

**C.** Tg supplementation does not alter mitochondrial respiration of PM-permeabilized H9 CHO cells. Oxygen consumption rate (OCR in pMole/min) was measured on a Seahorse XF-24 platform. For the "Tg" group (n=11 wells), wells received 250 nM Tg in the permeabilization buffer, while the “DMSO” group (n=10 wells) received same volume of DMSO as vehicle control. Both groups received 1 µM $Ca^{2+}$ in the form of $CaCl_2$ injected after the fifth measurement. Subsequently, both groups received sequential injections of carboxyatractyloside (**CAt,** 3 µM), FCCP (1 µM), and finally rotenone (**Rot**, 1 µM) plus antimycin A (**AA**, 10 µM). Data are represented as Mean ± SEM.

**D.** Tg supplementation does not alter mitochondrial respiration in PM-permeabilized H9 CHO cells. Data from panel (**C**) were used to calculate group averages after each inhibitor treatment, which are shown as bar graphs, with SEM represented by whiskers. Two-way ANOVA detected no effect of Tg at 250 nM, with a p-value of 0.97.

**E**. ATP production was estimated by measuring ATP content in the respiration buffer (50 µL out of 100 µL total volume) under the indicated EGTA and $Ca^{2+}$ concentrations, after 30 min of respiration in presence of 2 mM exogenous ADP at 37 ºC. Endogenous ATP in intact cells that were not permeabilized were included as an additional control for ATP level comparison. No significant difference was found between $Ca^{2+}$ groups below “*11100 nM* $Ca^{2+}$”. All groups received Tg treatment (250 nM) for 10 min before permeabilization.

**F**. Luminescence signals from the ATP standard curve is intended to provide an estimate of absolute amount of ATP contained in the respiration buffer.

**Figures S11**

**Raw data reflect meaningful changes of distinct imaging modules.**

For technical illustration purposes, original fluorescence intensities of the FRET, CFP and GFP signals are shown in response to DMSO (**A**), Tg (**B**), IAA (**C**) and oligomycin (**D**). The raw values were used to calculate the FRET-to-CFP ratios shown in **Figs. 3 H**& **I**.

**Figure S12**

**While mitochondrial matrix ATP levels remain unchanged in ER stressed H9 cells, upon OxPhos inhibition the cytosolic ATP/ADP ratios decrease more quickly than unstressed H9 cells.**

Mitochondrial ATP levels are not changed in un-induced H9 CHO cells (**A**) and SAHA-treated H9 cells to induce F8 expression (**B**). Cells were equally sensitive to 1 µM oligomycin. ATP levels were estimated by the *mtAT1.03* mitochondrial localized ATP reporter, through flow cytometry-based ratiometric measurements.

Cytosolic ATP/ADP ratios were monitored by the *PercevalHR* probe upon OxPhos inhibition by oligomycin plus rotenone (Oligo and Rot, respectively). ER stress induced by Tm (100 ng/mL x 18 hrs) was coupled with a faster decrease in the cytosolic ATP/ADP ratio upon OxPhos blockade. **C.** Cytosolic ATP/ADP ratio in ER stressed H9 CHO cells and their non-stressed counterparts is expressed as the percentage of that measured at 0 min time point for their respective groups. Round symbols represent cells that were not ER stressed, and square symbols represent ER stressed cells. Solid lines are cells treated with DMSO as vehicle control, while dotted lines are cells that received 1 μM oligomycin plus 2 μM rotenone. *Dunnett's* multiple comparisons test was subsequently performed for comparison between vehicle-treated versus OxPhos blockers-treated groups. **D.** Raw cytosolic ATP/ADP ratios from which the percentage values were derived in panel (**C**) is shown.

**Figures S13**

**Work flow for flow-cytometry based ATP level measurements in single cells.**

**Figures S14**

**A single clone of H9-D2 cells was engineered to facilitate flow-cytometry based ER ATP analysis.**

After ERAT4$^{N7Q}$ transfection and G418 selection, a single clone of H9 cells, H9-D2 cells, was further obtained to facilitate the flow-based FRET assay. (**A**-**B**). H9-D2 cells have unanimous ERAT4$^{N7Q}$ expression with reduced R-square of FRET vs. YFP, in comparison to the unselected H9 cell population. The *FRET ratio* vs. *YFP fluorescence* plots were generated by *FlowJO* software for comparison purposes. (n = 41701 events for **A** and 74224 for **B**, respectively). (**C**-**D**). When reporter positive cells (defined as YFP$^{hi}$ population) were gated out for analysis, data sphericity for FRET ratio was improved for H9-D2 cells (n = 8202 events for **C** and 73617 for **D**, respectively). The FRET ratio generated by H9-D2 cells allowed more accurate data analysis through *ANOVA*.

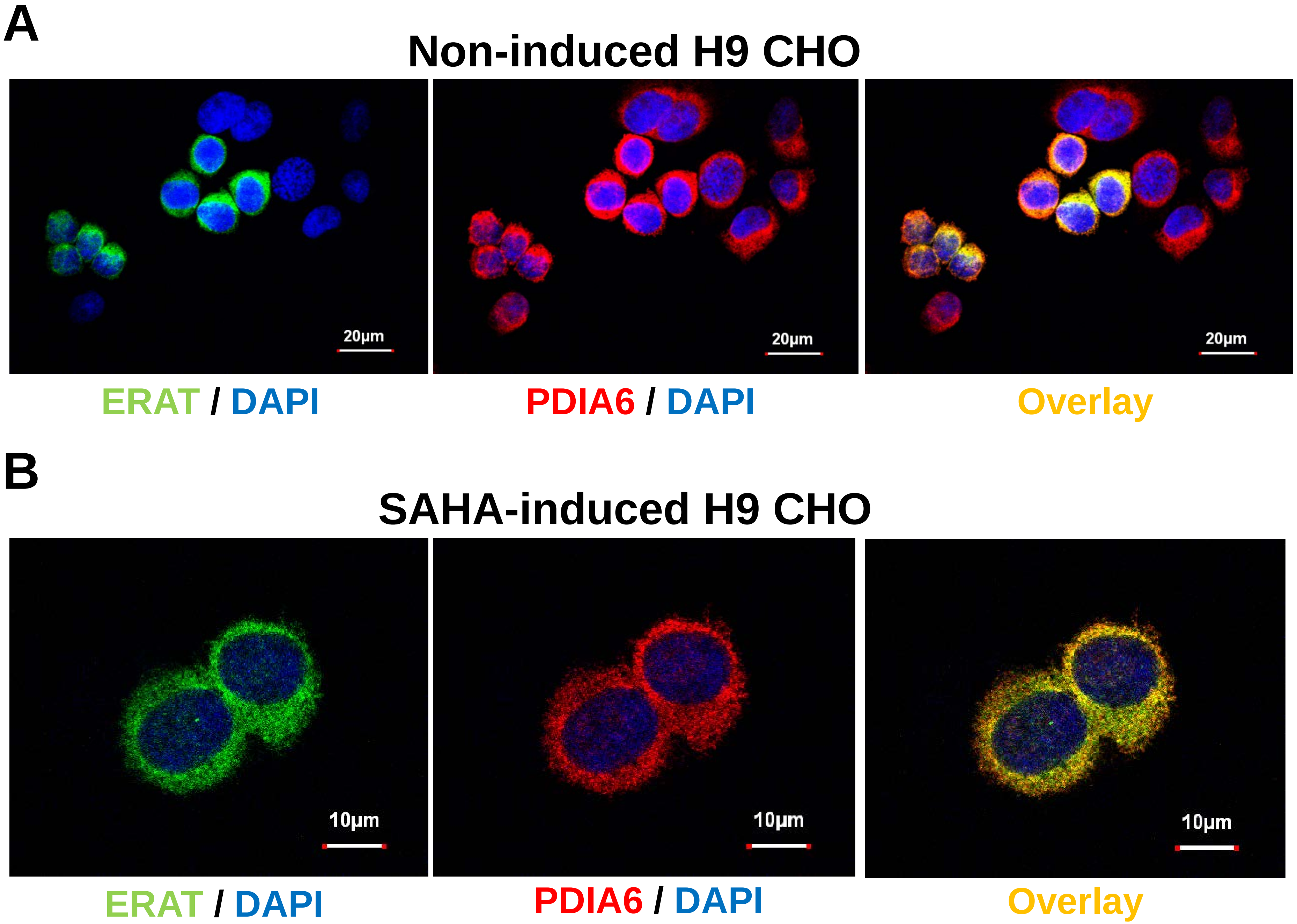
A
Non-induced H9 CHO
20μm
20μm
20μm
ERAT / DAPI
PDIA6 / DAPI
Overlay
B
SAHA-induced H9 CHO
10μm
10μm
10μm
ERAT / DAPI
PDIA6 / DAPI
Overlay

Yong et al., Sppl Figure S2

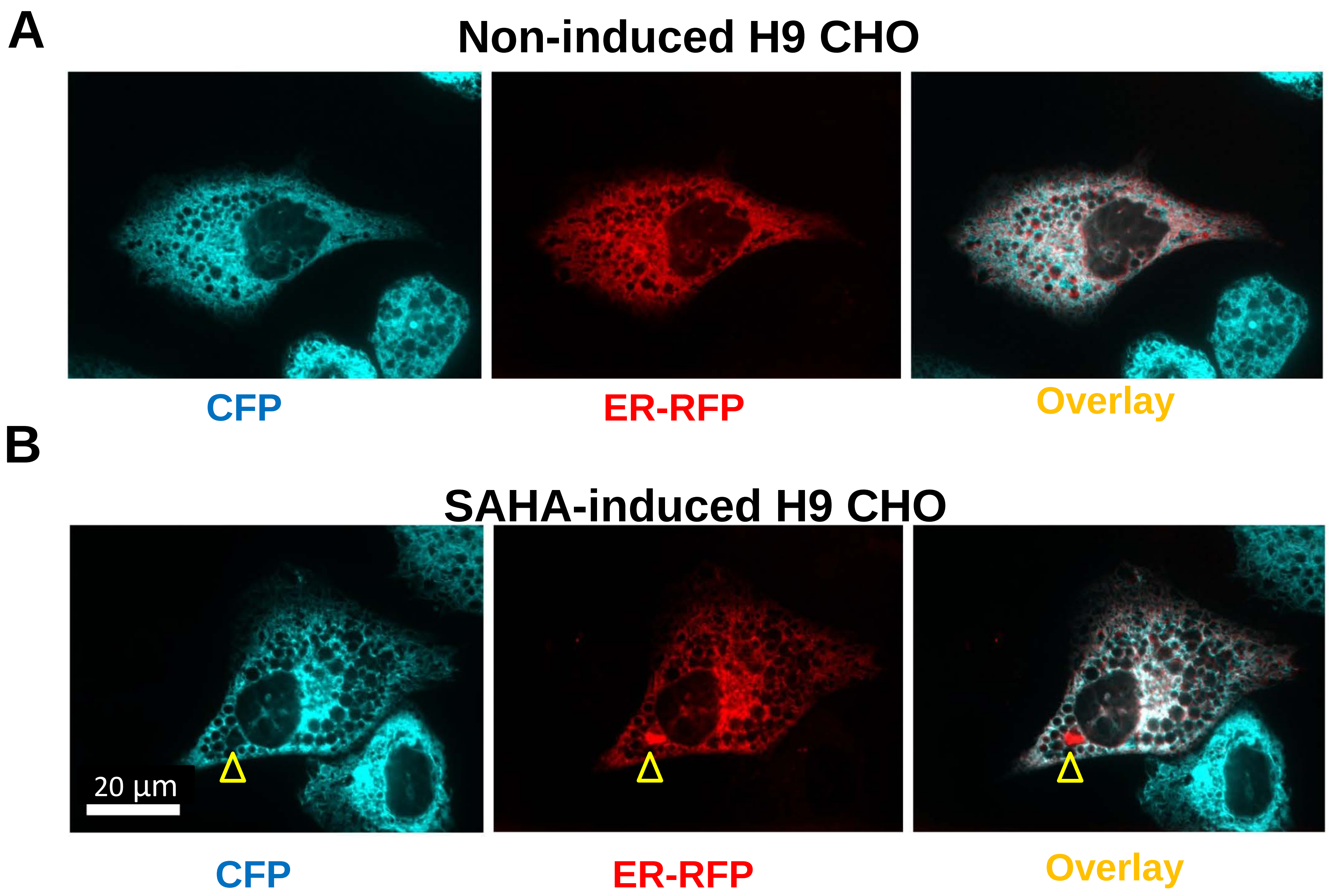

Yong et al., Sppl Figure S3

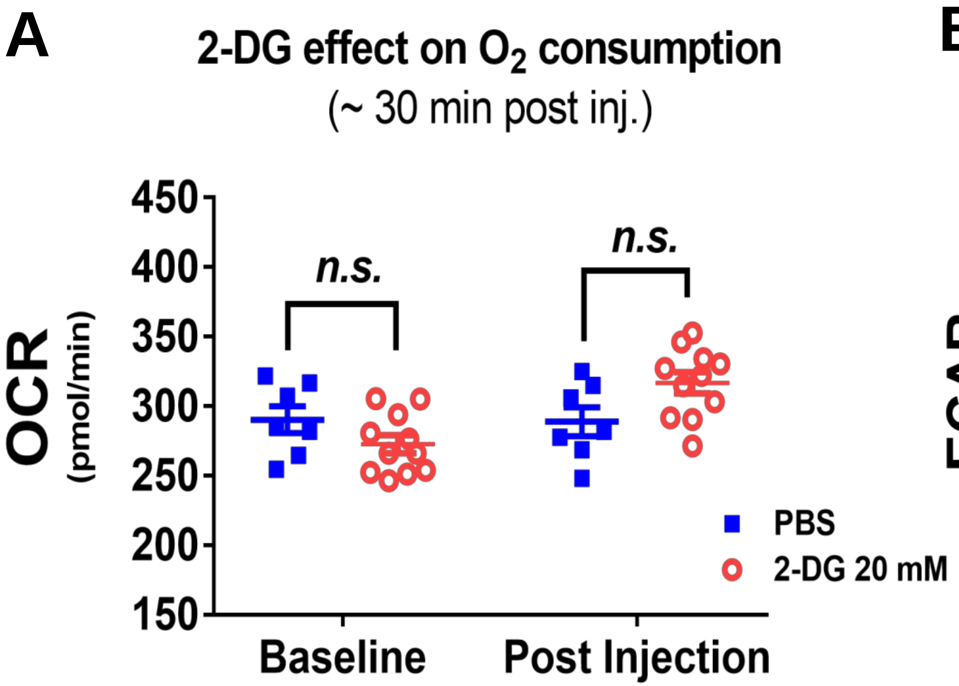


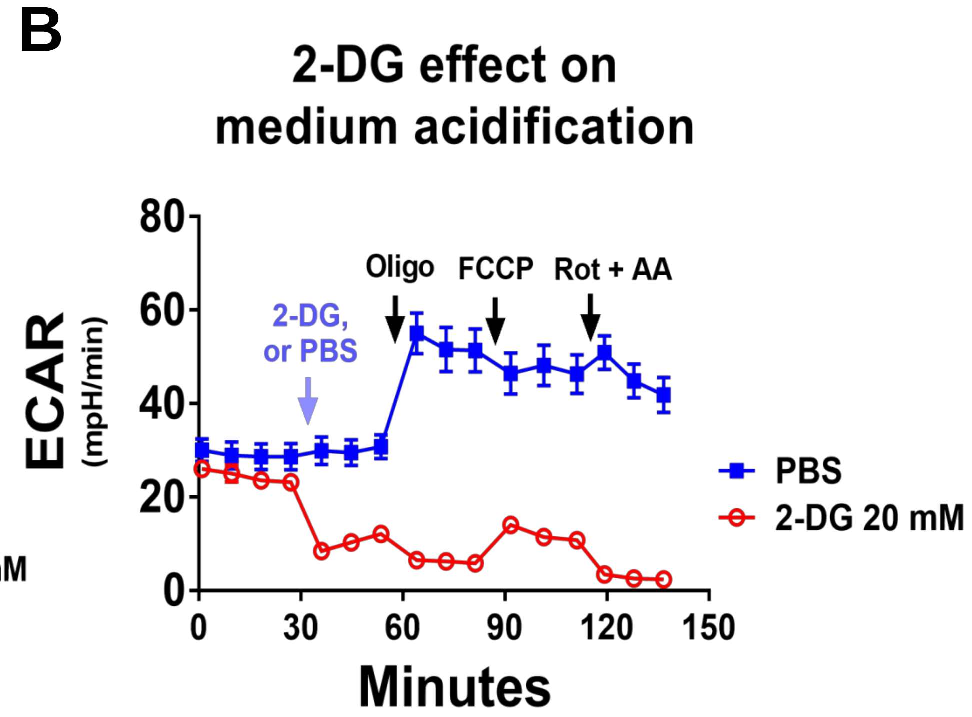

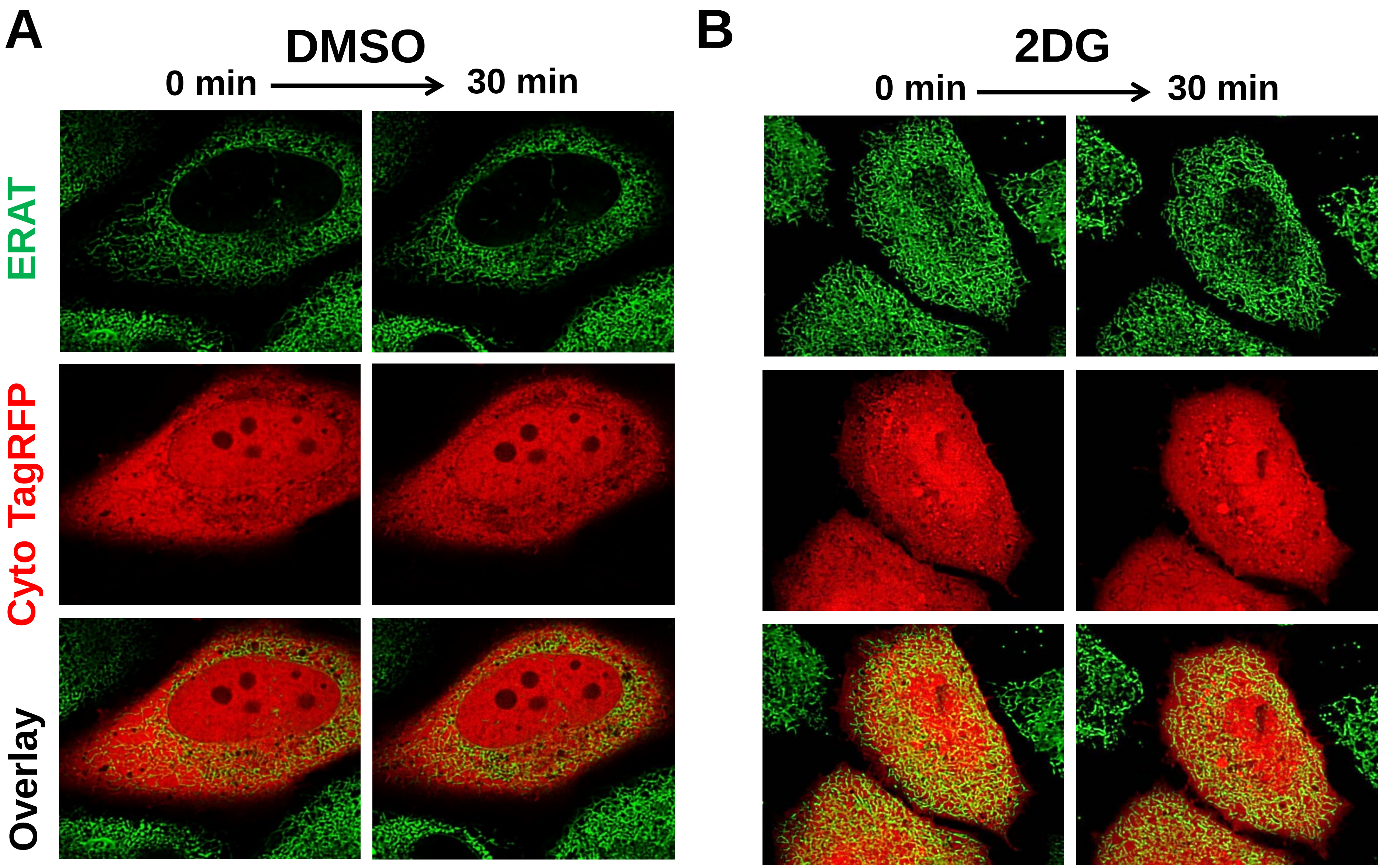
A
DMSO
0 min
30 min
ERAT
Cyto TagRFP
Overlay
B
2DG
0 min
30 min

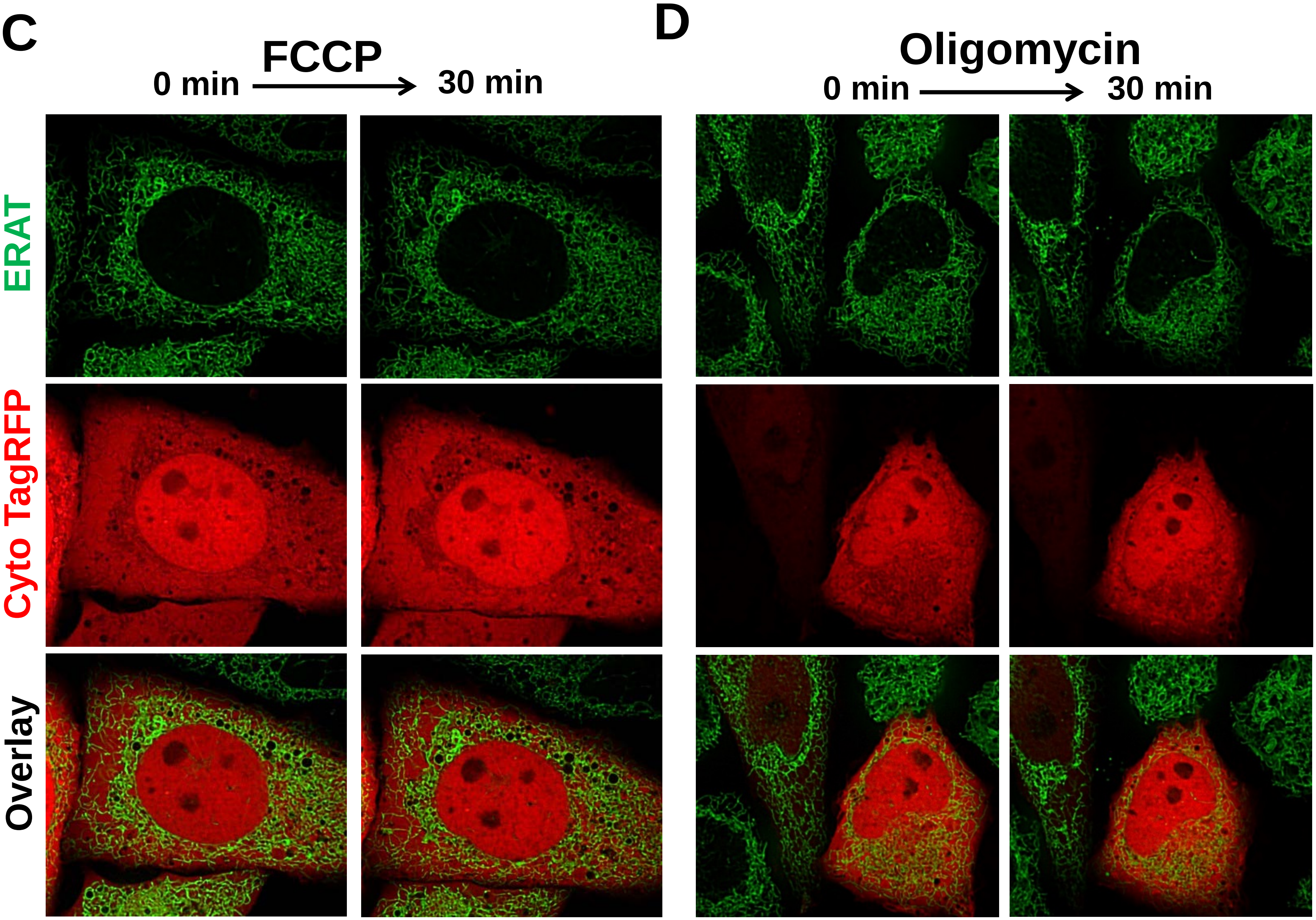
C
FCCP
0 min
30 min
ERAT
Cyto TagRFP
Overlay
D
Oligomycin
0 min
30 min

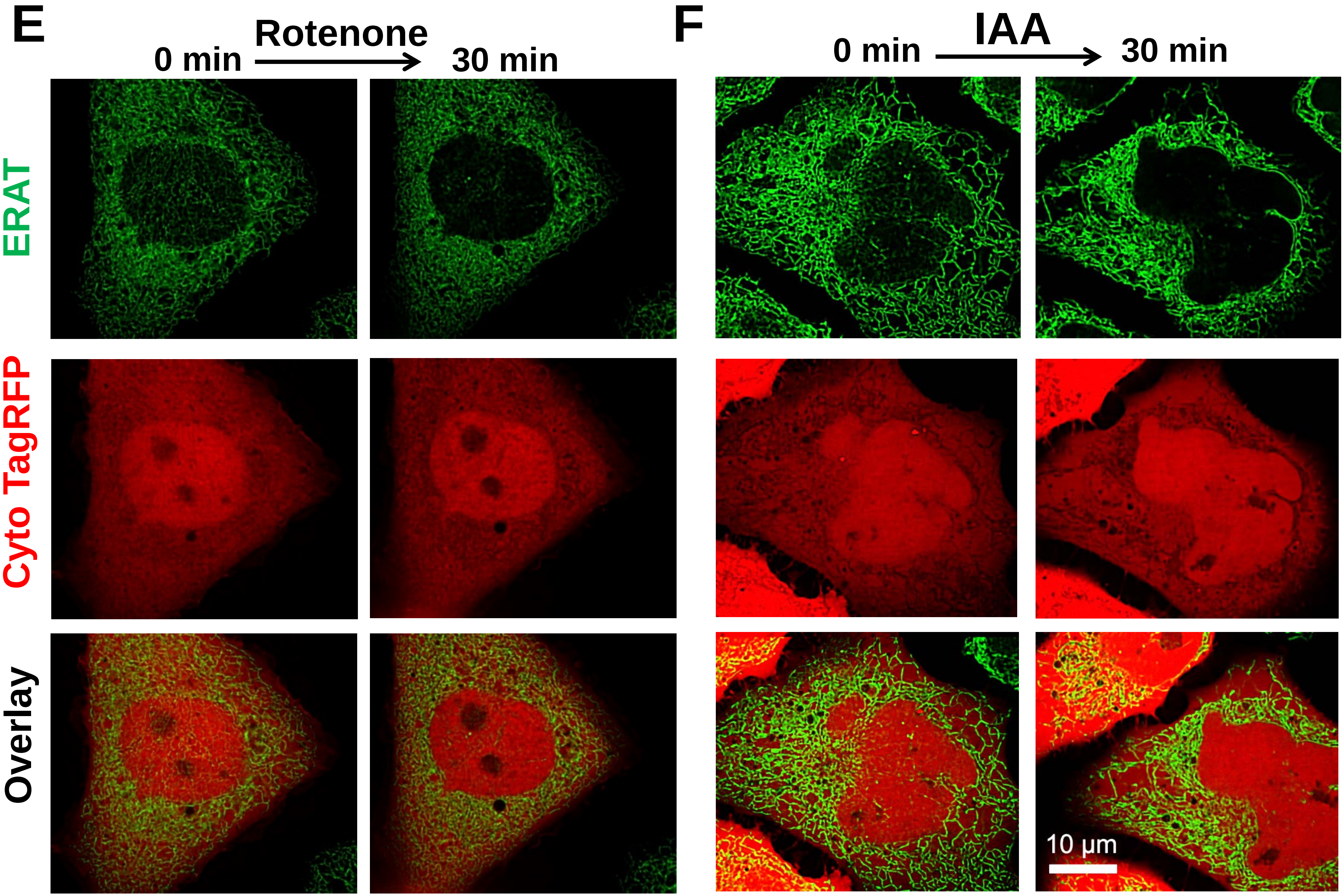
E
Rotenone
0 min
30 min
ERAT
Cyto TagRFP
Overlay
F
IAA
0 min
30 min
10 μm

G

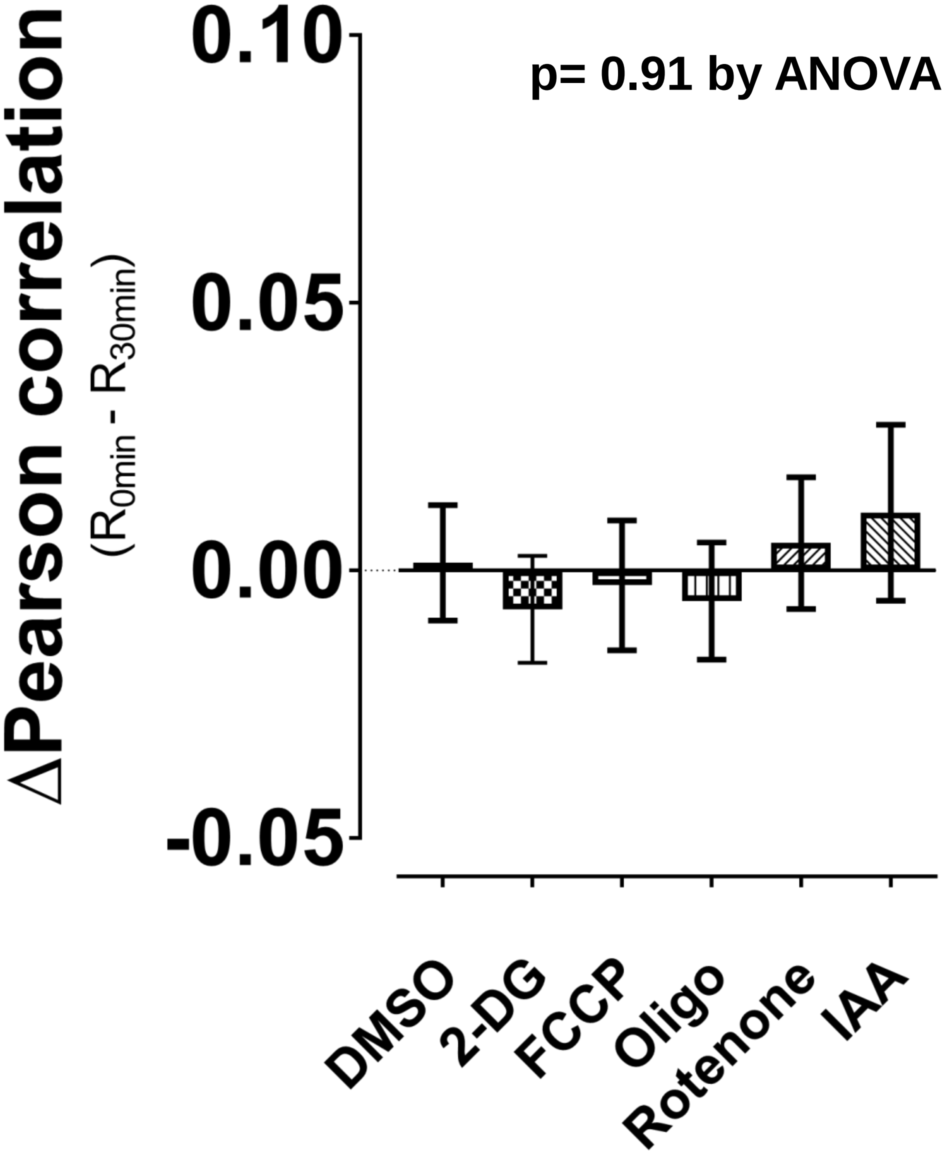
ΔPearson correlation
($R_{0min}$ - $R_{30min}$)
0.10
0.05
0.00
-0.05
p= 0.91 by ANOVA
DMSO
2-DG
FCCP
Oligo
Rotenone
IAA

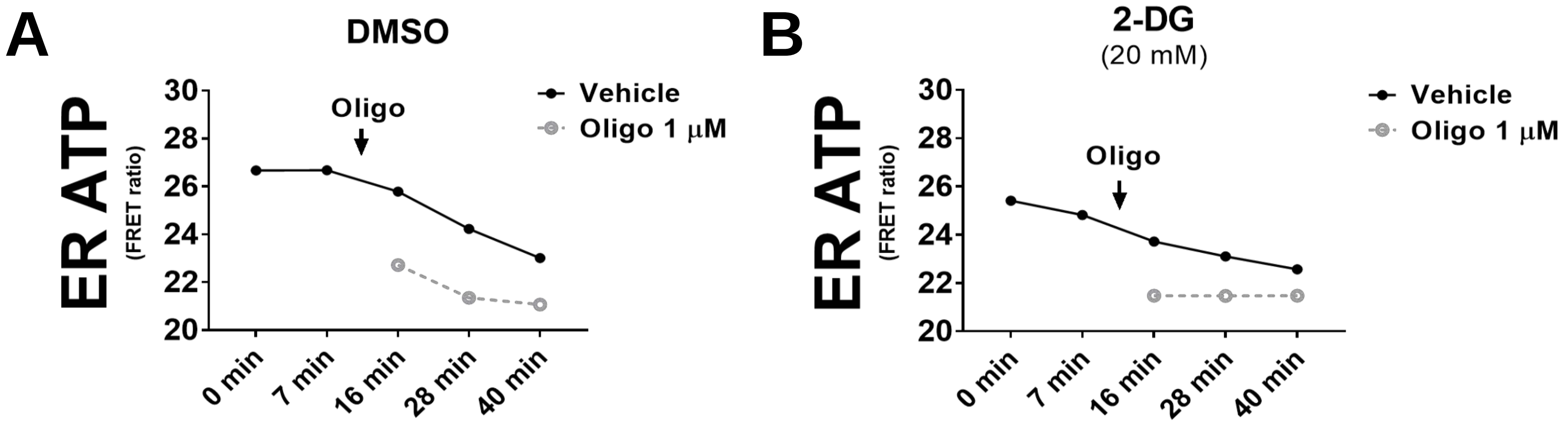
A
DMSO
ER ATP
(FRET ratio)
30
28
26
24
22
20
Oligo
Vehicle
Oligo 1 μM
0 min
7 min
16 min
28 min
40 min
B
2-DG
(20 mM)
ER ATP
(FRET ratio)
30
28
26
24
22
20
Oligo
Vehicle
Oligo 1 μM
0 min
7 min
16 min
28 min
40 min

Yong et al., Sppl Figure S6

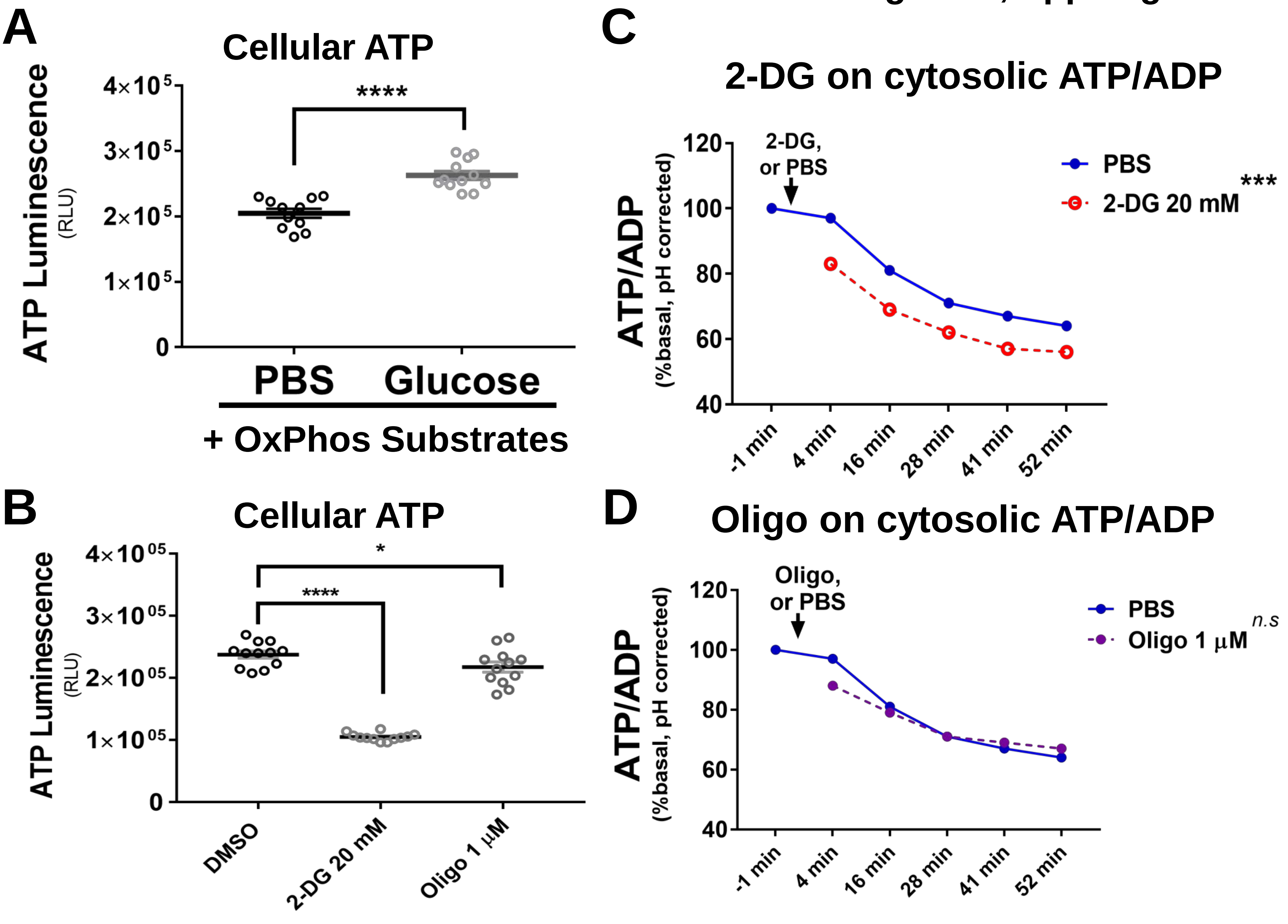

Yong et al., Sppl Figure S7

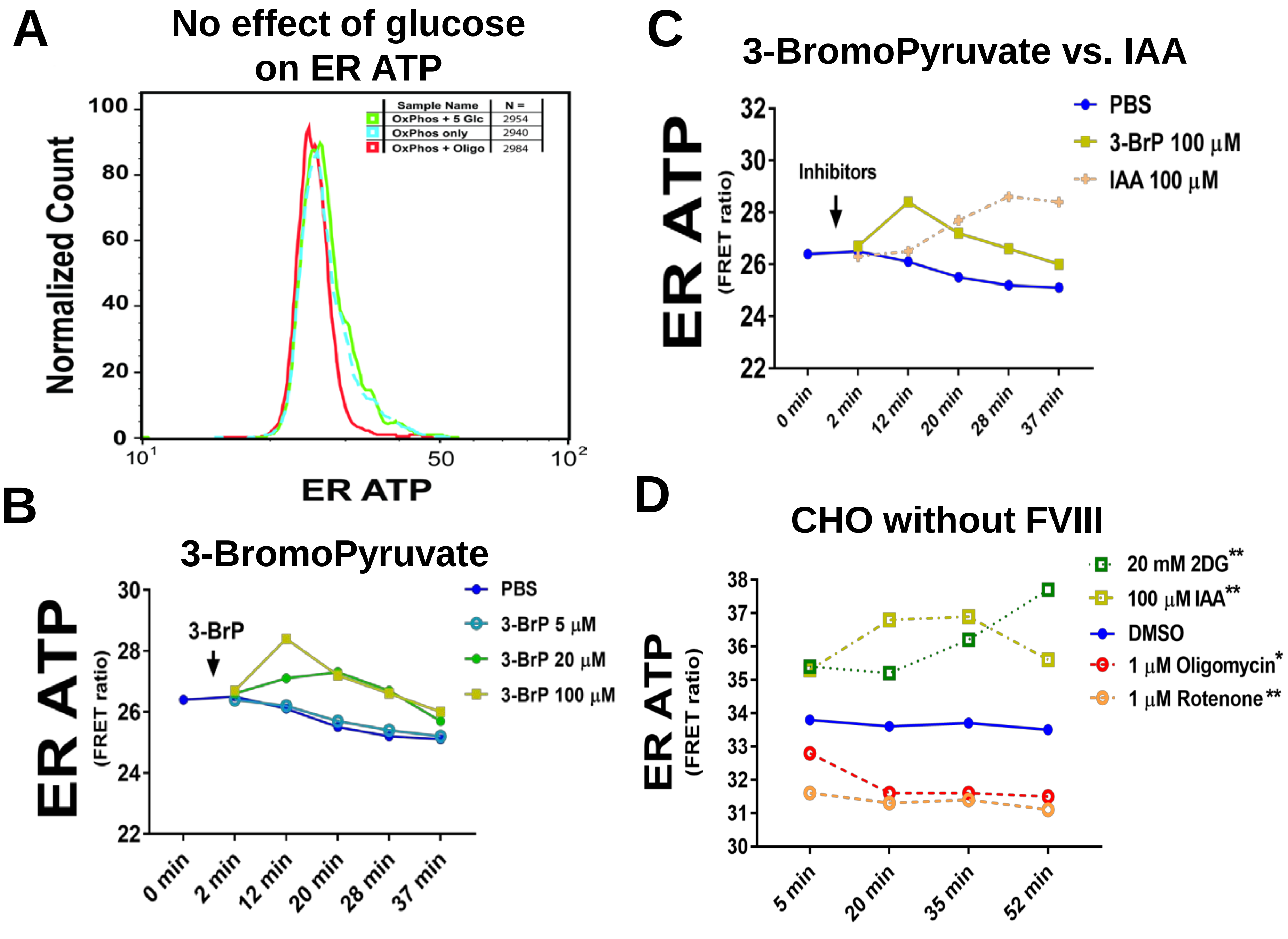

Yong et al., Sppl Figure S8

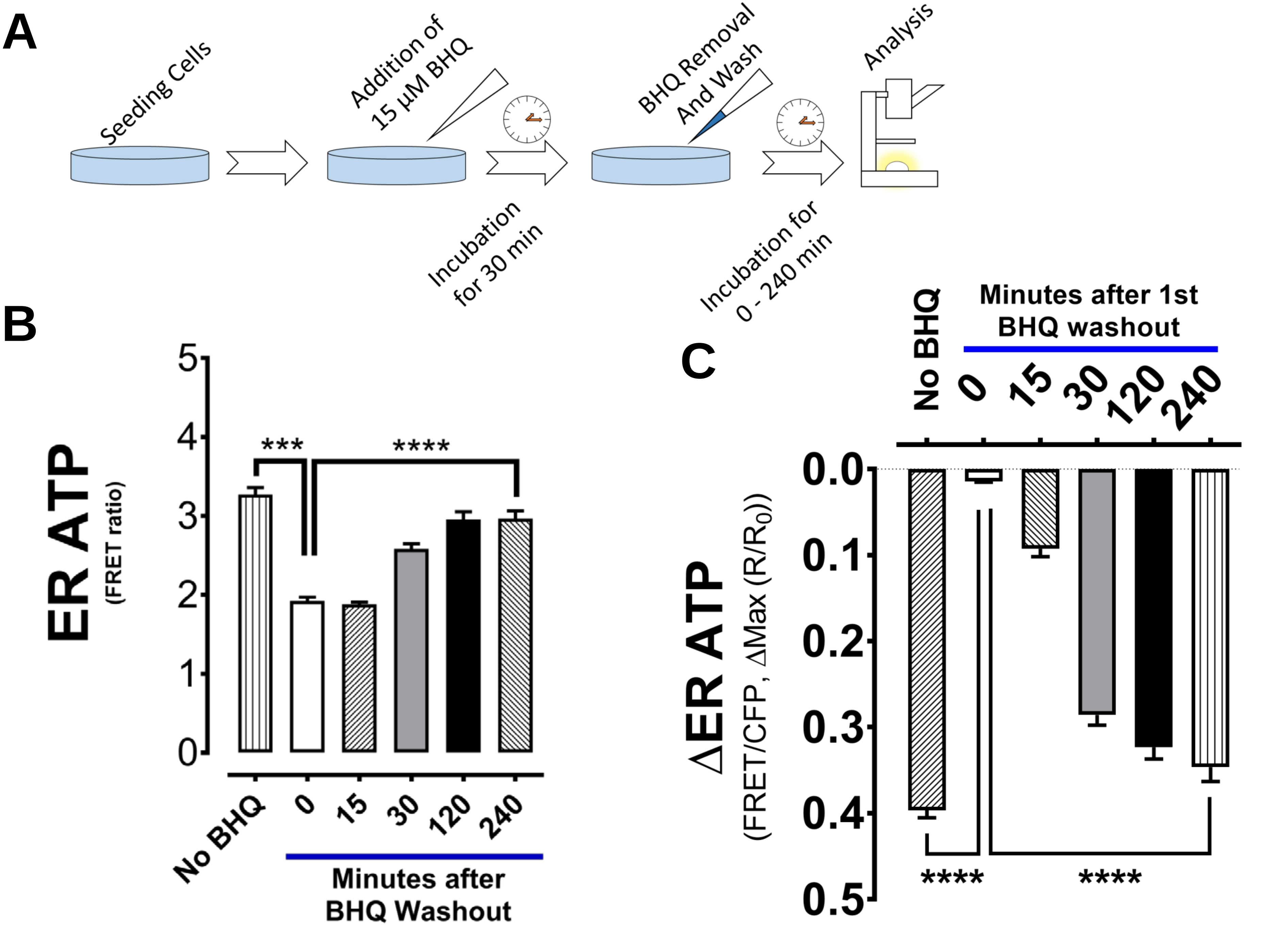

Yong et al., Sppl Figure S9

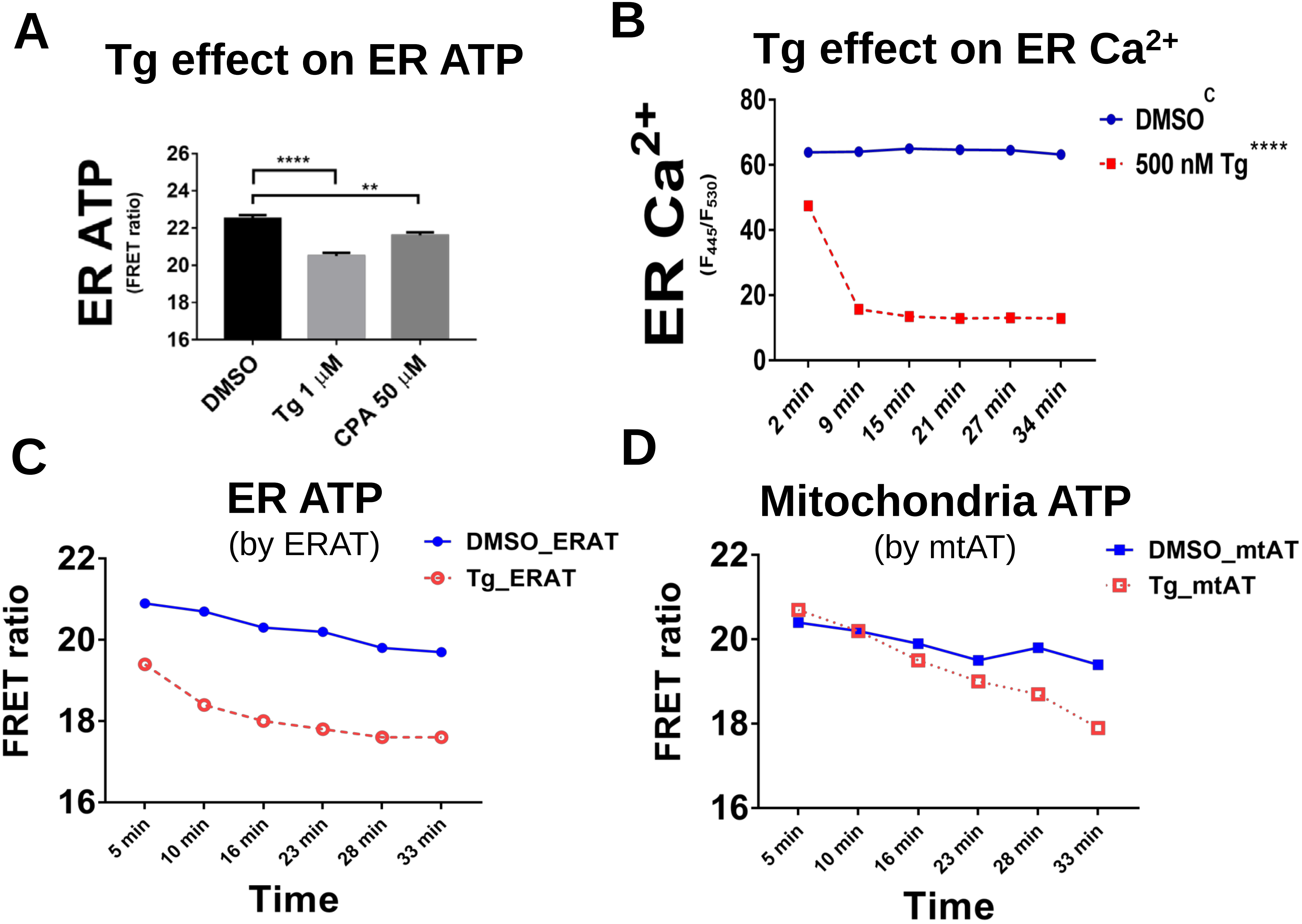

Yong et al., Sppl Figure S10

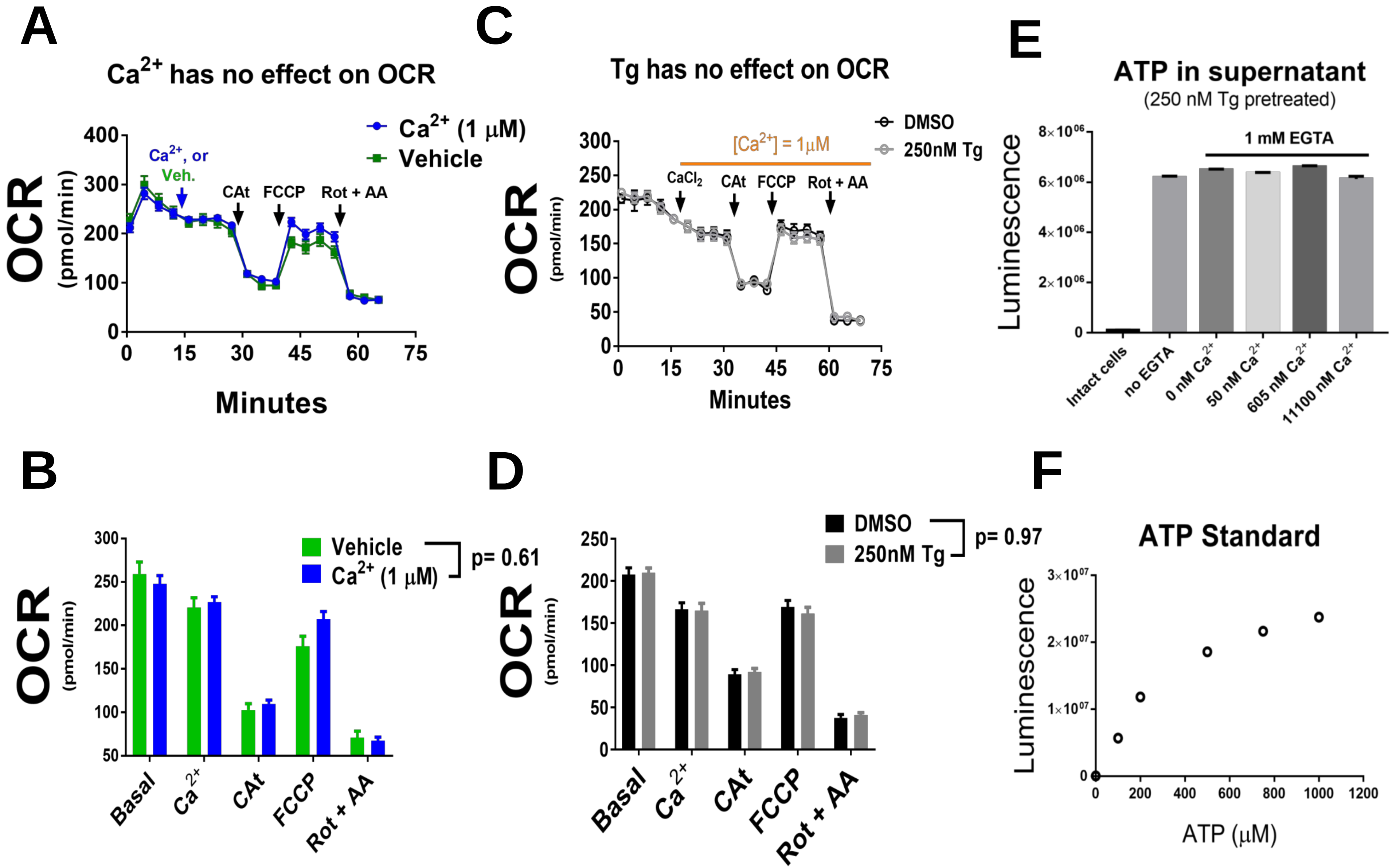

### Yong et al., Sppl Figure S11

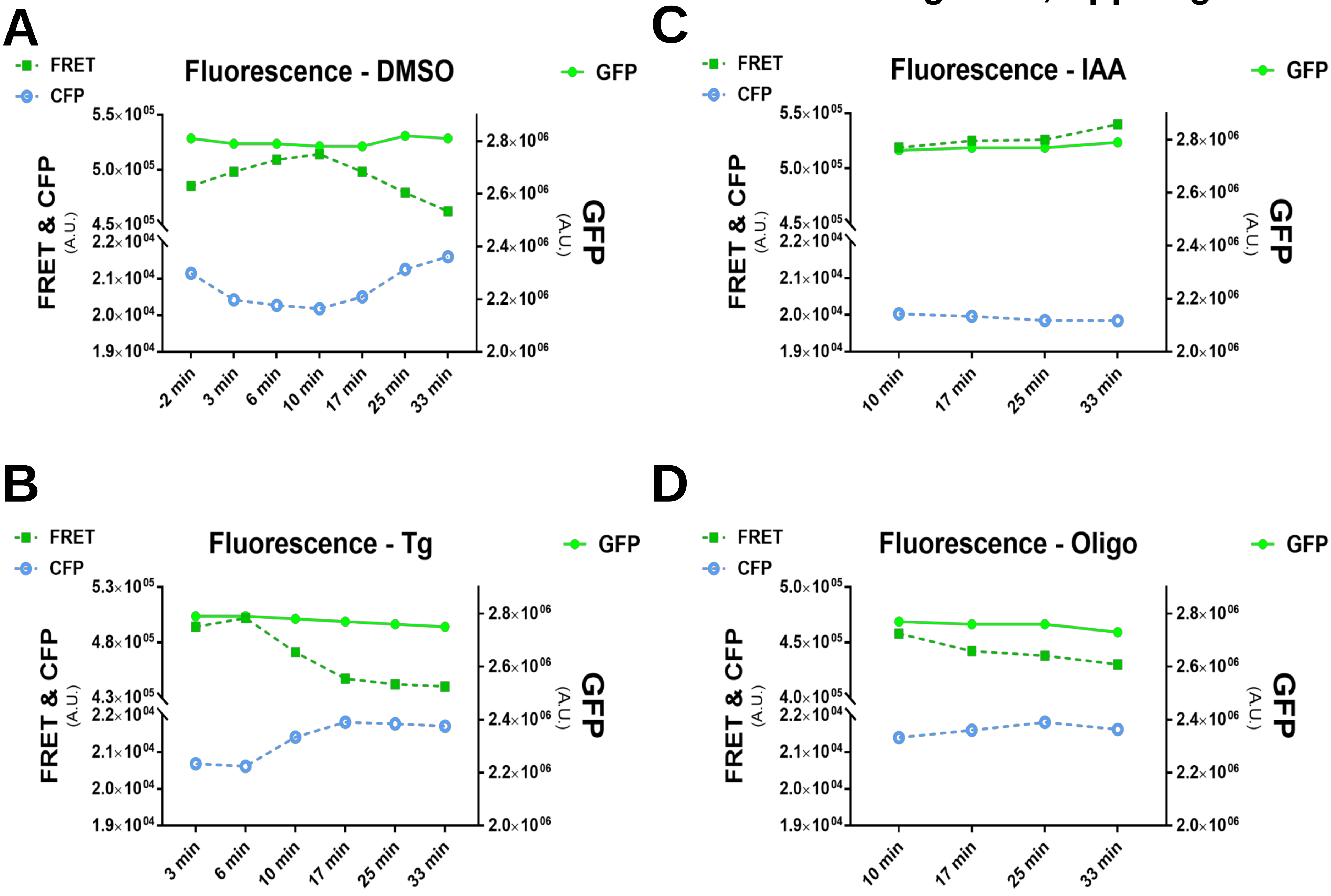

Yong et al., Sppl Figure S12

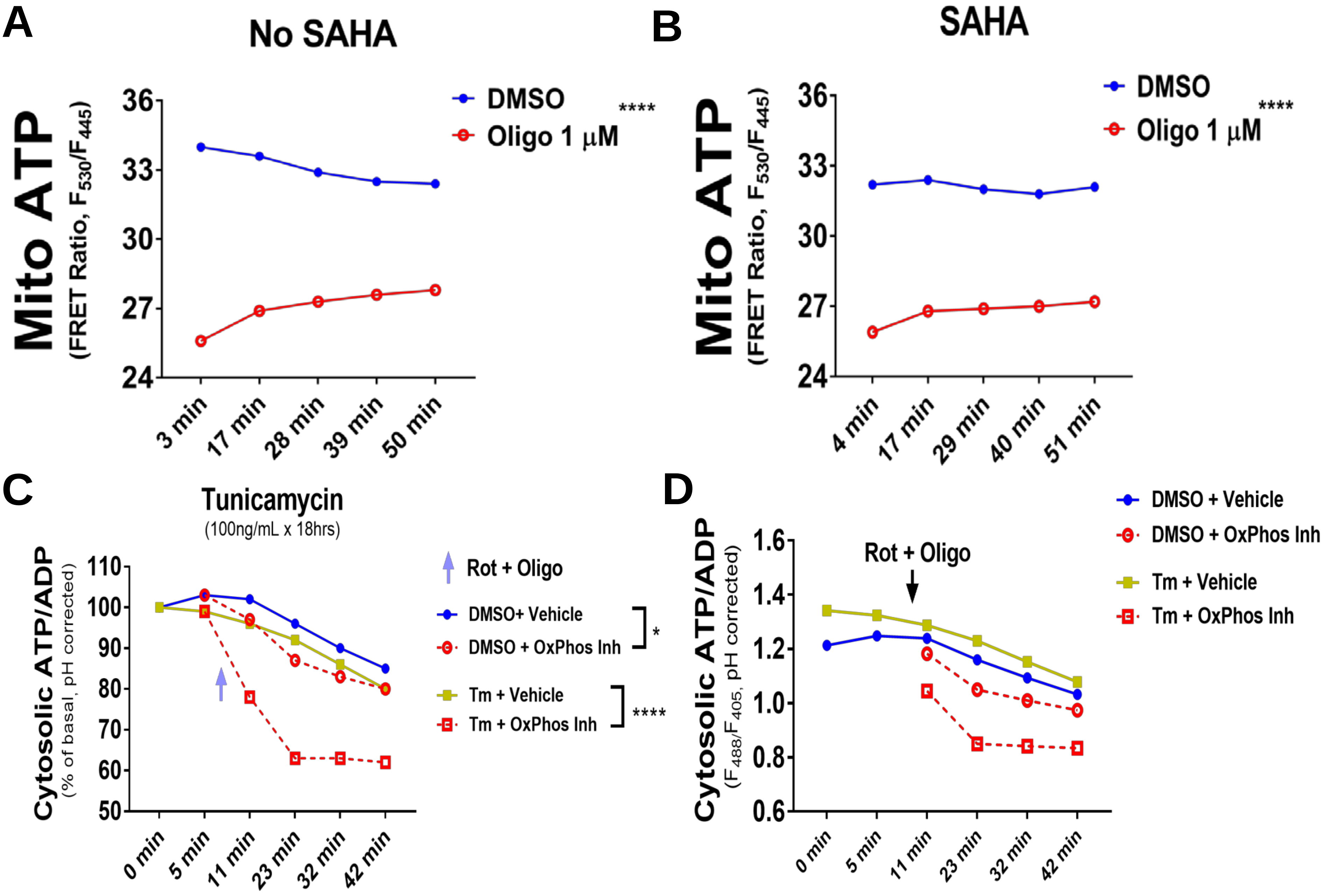

Yong et al., Sppl Figure S13

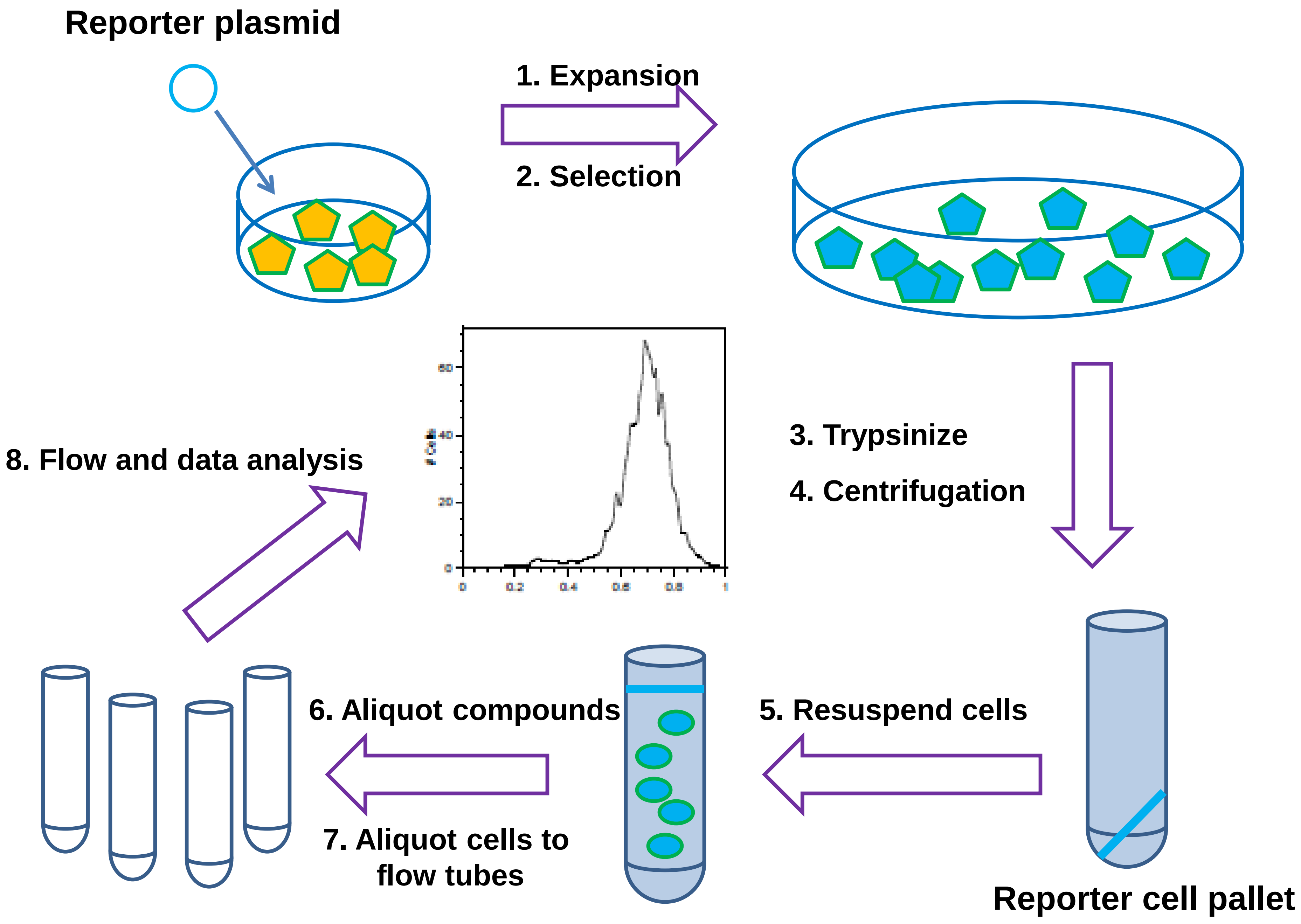

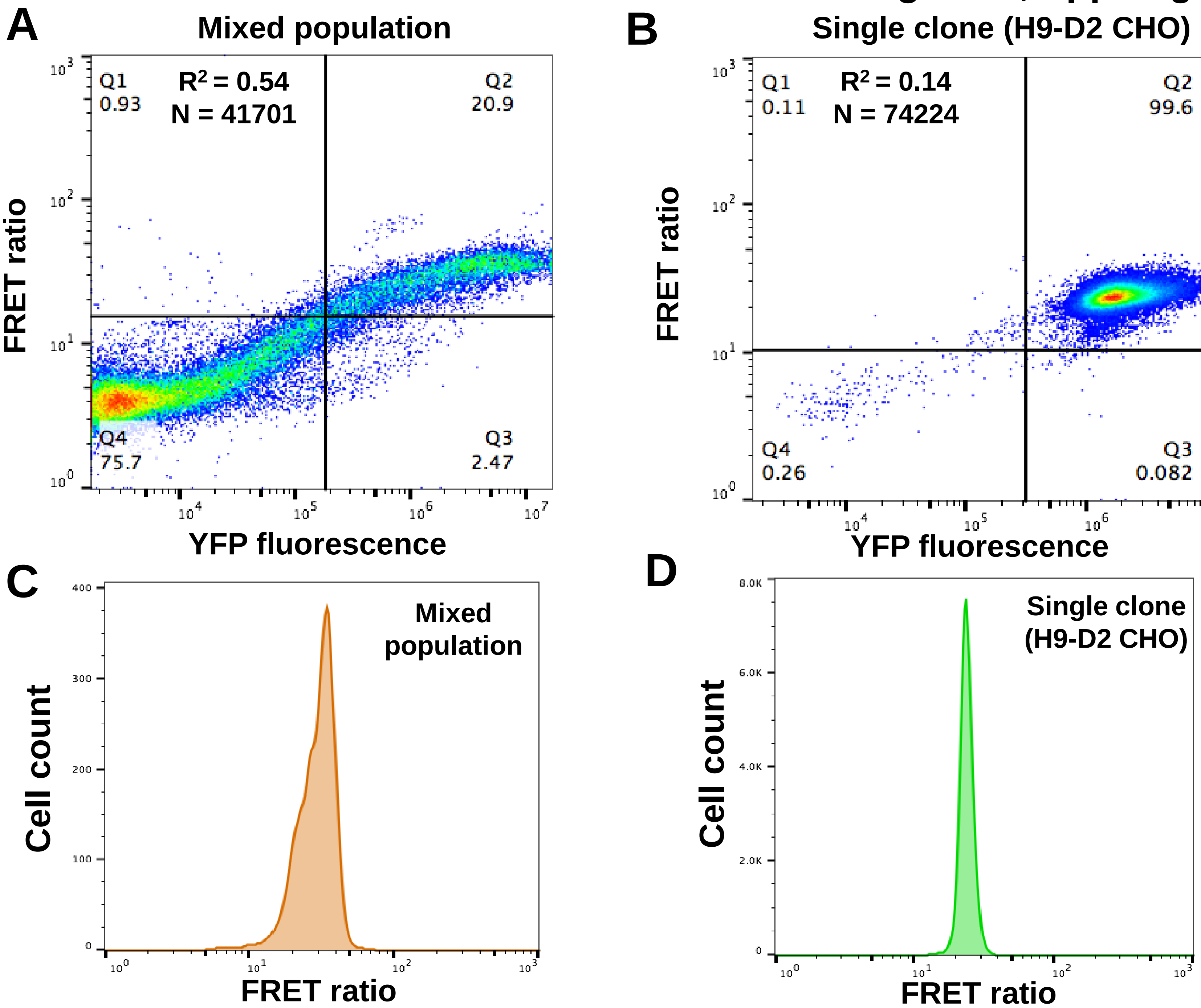
A
Mixed population
Q1
0.93
$R^2$ = 0.54
N = 41701
Q2
20.9
Q4
75.7
Q3
2.47
FRET ratio
YFP fluorescence
B
Single clone (H9-D2 CHO)
Q1
0.11
$R^2$ = 0.14
N = 74224
Q2
99.6
Q4
0.26
Q3
0.082
FRET ratio
YFP fluorescence
C
Mixed
population
Cell count
FRET ratio
D
Single clone
(H9-D2 CHO)
Cell count
FRET ratio
